# Protein Structure Inspired Drug Discovery

**DOI:** 10.1101/2024.05.17.594634

**Authors:** Fangfang Qiao, T. Andrew Binknowski, Irene Broughan, Weining Chen, Amarnath Natarajan, Gary E. Schiltz, Karl A. Scheidt, Wayne F. Anderson, Raymond Bergan

## Abstract

Drug discovery starts with known function, either of a compound or a protein, in-turn prompting investigations to probe 3D structure of the compound-protein interface. As protein structure determines function, we hypothesized that unique 3D structural motifs represent primary information denoting unique function that can drive discovery of novel agents. Using a physics-based protein structure analysis platform developed by us, designed to conduct computationally intensive analysis at supercomputing speeds, we probed a high-resolution protein x-ray crystallographic library developed by us. We selected 3D structural motifs whose function was not otherwise established, that offered environments supporting binding of drug-like chemicals and were present on proteins that were not established therapeutic targets. For each of eight potential binding pockets on six different proteins we accessed a 60 million compound library and used our analysis platform to evaluate binding. Using eight-day colony formation assays acquired compounds were screened for efficacy against human breast, prostate, colon and lung cancer cells and toxicity against human bone marrow stem cells. Compounds selectively inhibiting cancer growth segregated to two pockets on separate proteins. The compound, Dxr2-017, exhibited selective activity against human melanoma cells in the NCI-60 cell line screen, had an IC50 of 19 nM against human melanoma M14 cells in our eight-day assay, while over 2100-fold higher concentrations inhibited stem cells by less than 30%. We show that Dxr2-017 induces anoikis, a unique form of programmed cell death in need of targeted therapeutics. The predicted target protein for Dxr2-017 is expressed in bacteria, not in humans. This supports our strategy of focusing on unique 3D structural motifs. It is known that functionally important 3D structures are evolutionarily conserved. Here we demonstrate proof-of-concept that protein structure represents high value primary data to support discovery of novel therapeutics. This approach is widely applicable.

**Author summary:** We introduce the concept that protein 3D structure represents primary information which can support downstream investigations, in this instance leading to the discovery of novel anticancer therapeutics.

## Introduction

It is established that protein structure determines function. The formation of specific three-dimensional (3D) structures serves to mediate specific functions. However, the analysis of 3D structure has inherent limitations that relate to its complexity and cost, especially in the context of how one 3D structure may interact with a different 3D structure, as may by the case when one is considering how a protein may interact with drug.[1–4] For this reason, studies of structure are typically undertaken only after there is some evidence supporting potential functional relevance.

Prior evidence of function underlie structure-based drug design approaches.[5] While its central focus is understanding structure for the purpose of designing a drug that can interact with it, in reality the initial steps in this process are based on prior evidence that the target structure is considered to be biologically important.[6] Such evidence may stem from primary protein sequence and its association with function, function of the whole protein, or a host of other sources of information indicative of function. However, efforts that involve first considering 3D structure as a type of primary source of information are not well represented. We undertook a series of investigations to examine the notion that 3D structure represents an important source of primary information, that unique structure represents unique function that in turn can inform downstream application.

The process of drug discovery is well suited to such an application as therapeutic agents mediate their effects by binding to specific sites on proteins. Such sites, by definition, are functionally relevant, and the binding of a given therapeutic agent alters that function in a desirable manner. Target identification in drug discovery constitutes a central focus, and there are two general approaches: Targeted Drug Discovery (TDD) and Phenotype-based Drug Discovery (PDD). While TDD begins the drug discovery process with knowledge of the protein target, ideally its 3D structure, it stems from more primary evidence that supports the role of the target in disease.[7] An important limit to TDD relates to the complexity of disease and that a pre-identified single target may fail to address that complexity. A PDD approach addresses this limit, but is associated with the highly difficult task of deconvolution of complex biology and ultimate target identification.[8] Additional reasons for focusing on drug discovery relate its overall importance, its longstanding inherent limitations, and the need for new approaches. Longstanding limitations relate to failure rates of over 96% on going from initial discovery of an active agent in the lab to attainment of FDA approval, over 90% for drugs entering clinical trials, spiraling costs with current average estimates around $2 billion, and a development time frame of over 10 years.[6, 9–12]

To address the hypothesis that structure may serve as an important entry point into the process of drug discovery, we conducted a set of proof-of-concept investigations. We probed a library of experimentally determined protein crystal structures created by us using an integrated suite of analytics scripted to run on a supercomputer and designed by us to probe dynamic protein structure. We thereby identified 3D protein structures whose function was not otherwise characterized with the potential to bind drug-like compounds. Using this analytic suite, we probed for compounds predicted to bind, acquired them, and screened for efficacy and toxicity. One compound, Dxr2-017, inhibited human melanoma cell growth at low nM concentrations and exhibited little toxicity on human bone marrow stem cells at concentrations over 2100-fold higher. Dxr2-017 was shown to induce anoikis, a type of programmed cell death in need of effective therapeutics. These investigations provide proof-of-concept that protein structure represents a powerful untapped source of primary information that can be used across several applications, inclusive of drug discovery.

## Results

### Identification of unique structural motifs of high potential pharmacologic value

We are seeking to address the fundamental hypothesis that protein structure in-and-of-itself constitutes powerful information that can serve as a starting point from which to initiate the process of discovering novel therapeutics. This requires access to well annotated protein structure information and an ability to analyze it in an in depth and meaningful manner.

Using the Argonne National Laboratory Advanced Proton Source Synchrotron (APS), our group includes members of consortia that have constructed and maintained several beam lines, together allowing the acquisition of high-resolution protein structure information. In initiatives through the Midwest Center for Structural Genomics of the Protein Structure Initiative funded by the National Institute of General Medical Sciences (NIGMS) and the National Institute for Allergy and Infectious diseases (NIAID) funded Center for Structural Genomics of Infectious Diseases (CSGID), our group has used the APS as a platform to define a library of 3D protein structures.[13] Protein structures were determined using state-of-the-art equipment and facilities, were well annotated and accessible to the public through the Protein Data Bank (PDB). For the proof-of-concept goals of the current study, we considered protein structures deposited in the PDB through the CSGID initiative. The CSGID targets proteins that are from a priority list of bacterial pathogens. As one of many possible sampling approaches, we considered the first group of 220 deposited structures (**S1 Fig**).

From proteins in **S1 Fig** we used SurfaceScreen methodology to identify and consider potential binding sites for small molecules.[14–17] We have previously demonstrated that functional surfaces are the most highly conserved regions of a protein, that they exhibit strong ligand specificity, and that ligand binding preferences can be assessed in proteins lacking sequence similarity.[14, 18–21] SurfaceScreen allows for surface comparisons by decomposing them into global shape and local physiocochemical texture (**Fig 1**). In brief, the subcomponents of SurfaceScreen involve using ShapeSignature to characterize the global shape of a CASTp (Computed Atlas of Surface Topography of Proteins)[17] identified solvent accessible surface as a probability distribution,[22] and comparisons between the probability distributions of two surfaces conducted using Kolmogorov-Smirnov test methods.[23] Further, recognizing that biochemical function relies on the combination of shape and chemical compatibility, the latter, in the context of shape, is considered with SurfaceAlign. SurfaceAlign compares coordinate combination sets of two surfaces, formed by the different chemical properties of its constituent amino acid groups, through decomposition of single value coordinates to identify the least square rotational matric, translation vector and root mean square distance (RMSD),[24] and takes into consideration both coordinate and orientation RMSD.[20] Statistical methods incorporate random surface alignments, global surface volume overlap Tanimoto coefficient, and lead to the generation of a composite SurfaceScreen Score.[20, 25–28]

**Fig 1.**
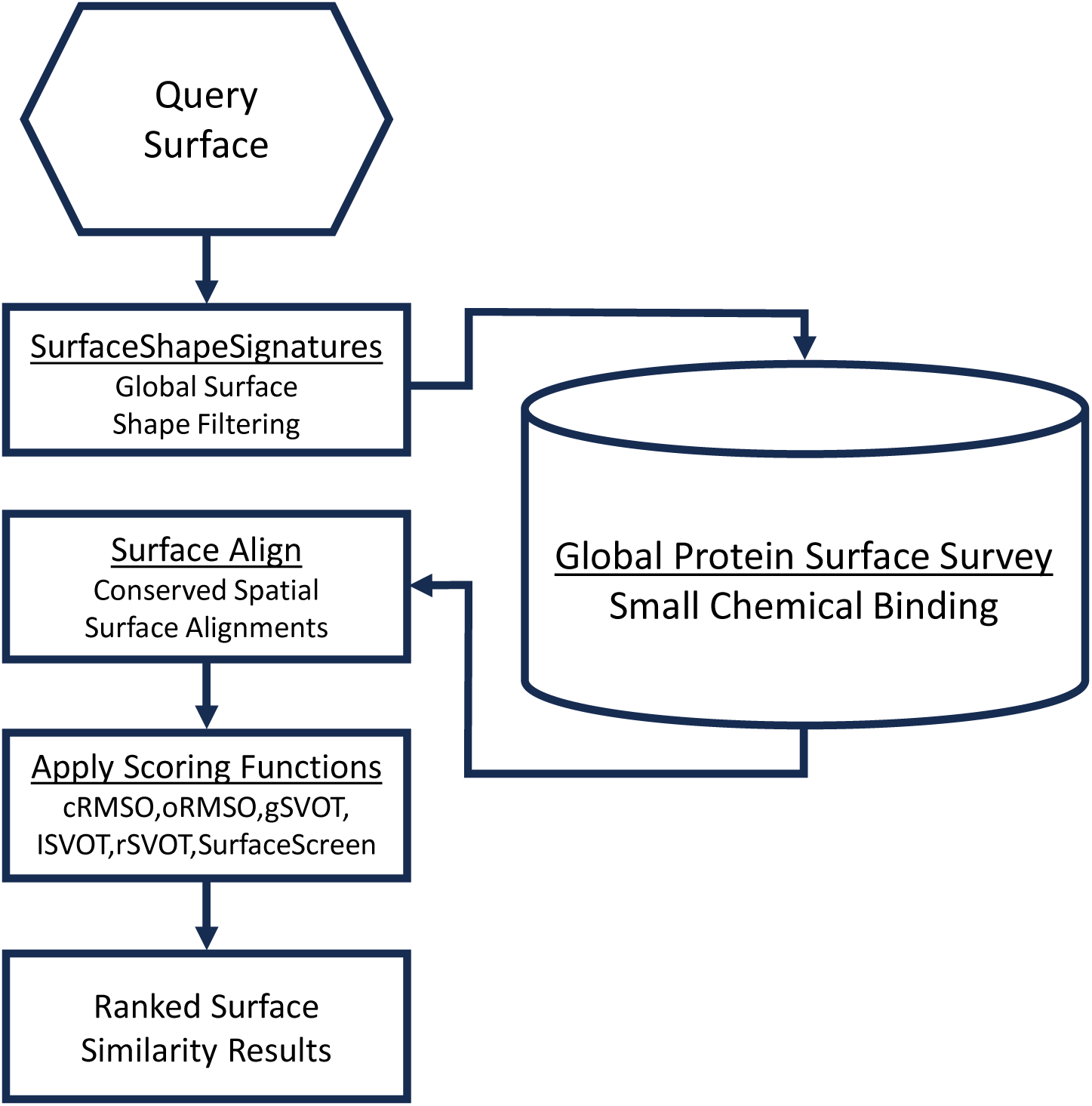
Schema for identification and characterization of 3D protein structures with the potential to bind therapeutically active small molecules.

We then sought to identify pockets on individual proteins that had the potential to serve as biologically important functional sites and that also had the potential to serve as target sites for small molecule therapeutics. We first sought pockets of 500 Å^2^, a value based on lower limit default settings with docking programs in common use.[29] Further, we sought the presence of at least two polar residues within a pocket. A value derived by considering that of commercially available drugs, from the Comprehensive Medicinal Chemistry (CMC) database, most side chains were polar, and the modal number of side chains was two.[30] We gave priority to pockets with a single mouth, based on findings that pockets for drug like ligands are almost always one mouthed.[31]

In making final selection of sites for further investigation we considered several additional factors. We prioritized sites whose function was not otherwise characterized. We sought sites exhibiting conservation of structure. During evolution highly conserved proteins are often required for basic cellular function, stability and folding kinetics.[32–39] Functional sites, such as enzyme active sites, are associated with the convergence of spatially conserved amino acid residues. Such conserved functional surfaces exhibit strong specificity and selectivity for compatible ligands.[14, 18–20] When residues that are conserved across species in a large sequence family come together in space, it indicates an evolutionary bias towards important functionality. This phylogenetic conservation of structure and function supports the hypothesis that important structural elements identified in bacterial proteins have functionally relevant counterparts in human cells. Surfaces with higher degrees of conservation were given higher priority, with those conserved into eukaryotes given the highest priority. Another factor we considered was the nature of the protein on which the pocket is located. We gave higher priority to proteins that, to our knowledge, did not have cognate small molecule modulators of their function that were advancing into clinical trials.

Taking the above factors into consideration we selected eight potential binding clefts on six different proteins for further investigation (**Fig 2**). Two proteins are expressed in bacteria through humans: 1. hypoxanthine-guanine phosphoribosyltransferase (HGPRT). This enzyme catalyzes the conversion of hypoxanthine or guanine to inosine or guanosine monophosphate, and is important in generating purine nucleotides through the purine salvage pathway.[40] Prior reports describe targeting HGPRT with tailor synthesized nucleoside analogs,[41] as well as by pentamidine, 1,3-dinitroadamantane, acyclovir and acyclovir analogs.[42, 43] The focus of these reports is on treatment of parasitic diseases, and higher activity is reported in HGPRT from parasites compared to that from human. 2. dihydrofolate synthase (FolC). There are two clefts on this protein, which we designate as FolC1 and FolC2. FolC catalyzes two reactions: its dihydrofolate synthetase activity catalyzes the addition of L-glutamate to dihydropteroate (7,8-dihydropteroate), forming dihydrofolate (7,8-dihydrofolate monoglutamate) and its folylpolyglutamate synthetase activity catalyzes subsequent additions of L-glutamate to tetrahydrofolate, forming folylpolyglutamate derivatives.[44] There are many inhibitors of other enzymes in the folate pathway in widespread clinical use, but those that target FolC are not among them. While FolC is expressed in humans, its dihydrofolate synthetase activity is not present in humans, and efforts to therapeutically target FolC appear to be limited, primarily focused on parasites, especially *P. falciparum*.[45]

**Fig 2.**
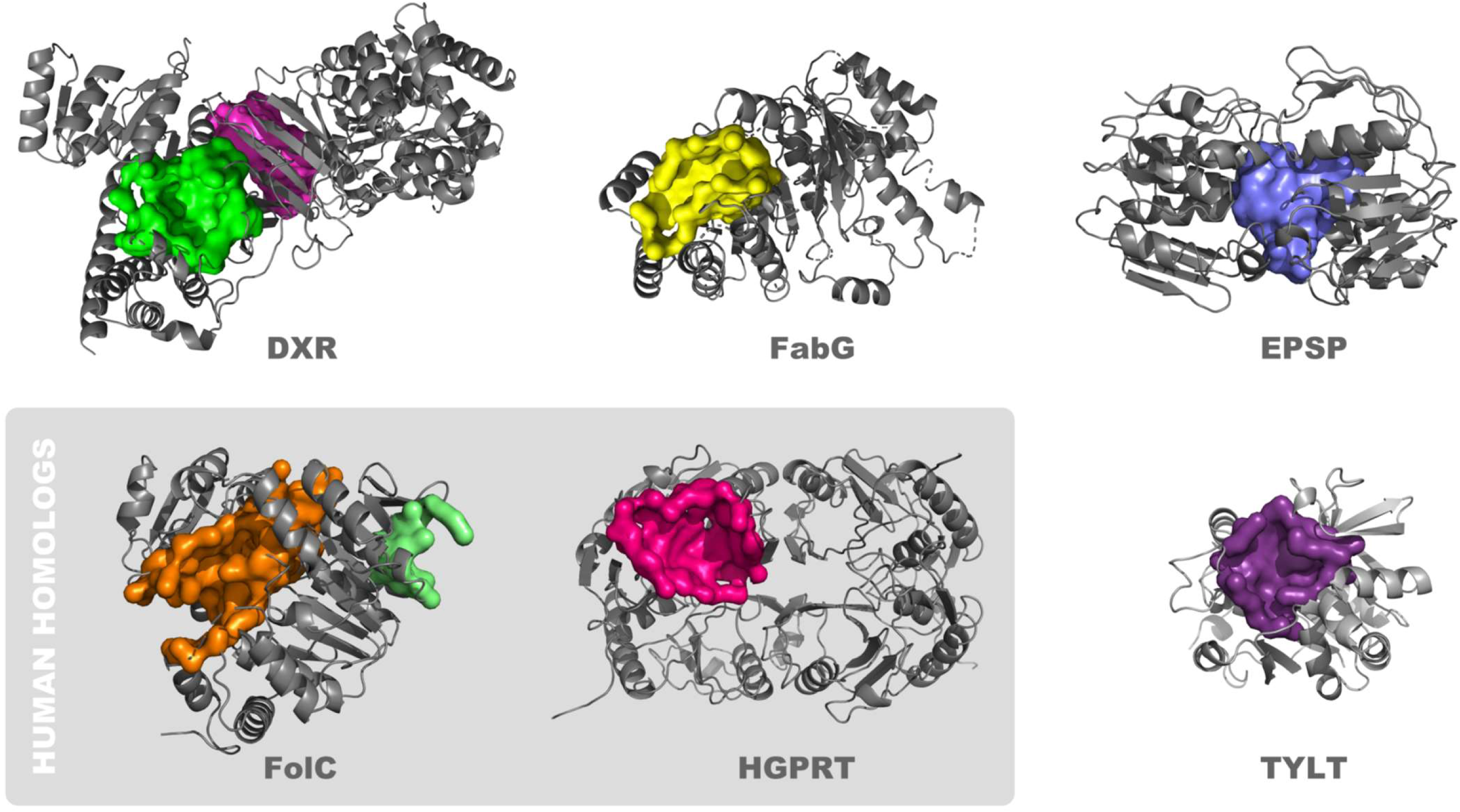
Proteins that contain pockets structurally suited to binding drug-like small molecules. The surfaces of potential binding pockets are depicted, as are ribbon structures of proximal portions of the protein. The proteins are: 1-deoxy-D-xylulose 5-phosphate reductoisomerase (Dxr; Dxr1 [magenta] and Dxr2 [green] binding pockets), ß-ketoacyl acyl carrier protein reductase (FabG), 3-phosphoshikimate 1-carboxyvinyltransferase (EPSP synthase), dihydrofolate synthase (FolC; FolC1 [orange] and FolC2 [light green] binding pockets), hypoxanthine-guanine phosphoribosyltransferase (HGPRT) and glucose-1-Phosphate Thymidylyltransferase (TYLT).

The other four proteins are primarily expressed in bacteria, and not in humans. 3. 1-deoxy-D-xylulose 5-phosphate reductoisomerase (Dxr), also known as DXP reductoisomerase. There are two pockets on this protein (Dxr1 and Dxr2). Dxr catalyzes the NADP-dependent rearrangement and reduction of 1-deoxy-D-xylulose-5-phosphate (DXP) to 2-C-methyl-D-erythritol 4-phosphate.[46] It is involved in the methylerythritol phosphate pathway and its synthesis of isopentenyl compounds. Several compounds have been shown to inhibit Dxr, and their use is geared toward treatment of bacterial and parasitic diseases, inclusive of tuberculosis and malaria.[47] 4. ß-ketoacyl acyl carrier protein reductase (FabG), also known as 3-oxoacyl-[acyl-carrier-protein] reductase. FabG catalyzes reduction of beta-ketoacyl-acyl carrier (ACP) protein substrates by NADPH to beta-hydroxyacyl-ACP products, can also catalyze reduction of acetoacetyl-CoA, albeit less efficiently than paralog proteins, and appears to play a role in fatty acid synthesis.[48, 49] Initially characterized in *E. coli*, FabG1-4 orthologs are present in *M. tuberculosis*,[50] and some have been shown to be inhibited by isoniazid,[51] which is widely used in the treatment of tuberculosis. 5. Glucose-1-Phosphate Thymidylyltransferase (TYLT). TYLT catalyzes condensation of substrates glucose-1 phosphate and deoxy-thymidine triphosphate (dTTP), producing pyrophosphate and dTDP-D-glucose,[52] the first step in L-rhamnose synthesis, which in turn is a cell wall component of bacteria, inclusive of *M. tuberculosis* and *P. aeruginosa*. Thymine analogs developed against TYLT were surprisingly found to act through binding of an allosteric site,[53] where current discovery efforts are focused,[54] and not the active site, which is the pocket of interest in the current study. 6. 3-phosphoshikimate 1-carboxyvinyltransferase (EPSP synthase). EPSP catalyzes the condensation of 3-phosphoshikimate + phosphoenolpyruvate, producing 5-O-(1-carboxyvinyl)-3-phosphoshikimate + phosphate.[55] This constitutes an essential step in the shikimate pathway,[56] present in bacteria through plant cells, but not animal cells, is used in the synthesis of folates and aromatic amino acids, and is targeted by the widely used herbicide, glyphosate.[57]

### Identification of small molecule ligands with the potential to interact with binding pockets

We next sought to identify small molecule ligands that have the potential to interact with identified pockets using our computational pipeline for docking and scoring (**Fig 3**).^19-21,23-26^ Docking simulations were against a library of 60 million compounds optimized for use in this pipeline and represent a subset of the ZINC compound library.[58] The library was first filtered for properties that have been associated with oral FDA approved small molecule therapeutics, inclusive of molecular weight between 160-480 g/mol and total polar surface area <140 square Angstroms (i.e., <5 and <10 hydrogen-bond donors and acceptors, respectfully).[59] These properties play a central role in defining a compound’s Adsorption, Distribution, Metabolism, and Excretion (ADME) characteristics, and thus desirable pharmacokinetic profile.[59] Using SurfaceScreen methodology, our pipeline initially incorporated AutoDock[60] and DOCK[61] docking applications, together providing a broad coverage of protein-small molecule docking strategies, including accounting for protein residue and compound flexibility. Top ranking initial docking poses are then ‘funneled’ into increasingly more sophisticated and computationally complex approximations of free energy of interaction using the MM-GBSA[62] (molecular mechanics-generalized Born surface area) and FEP/MD-GCMC[63] (molecular dynamics free energy perturbation-grand canonical Monte Carlo) methods, with rescoring of poses conducted in a hierarchical fashion. In this manner, we created a ranked list of top 100 compounds for each pocket (**S2 Fig**).

**Fig 3.**
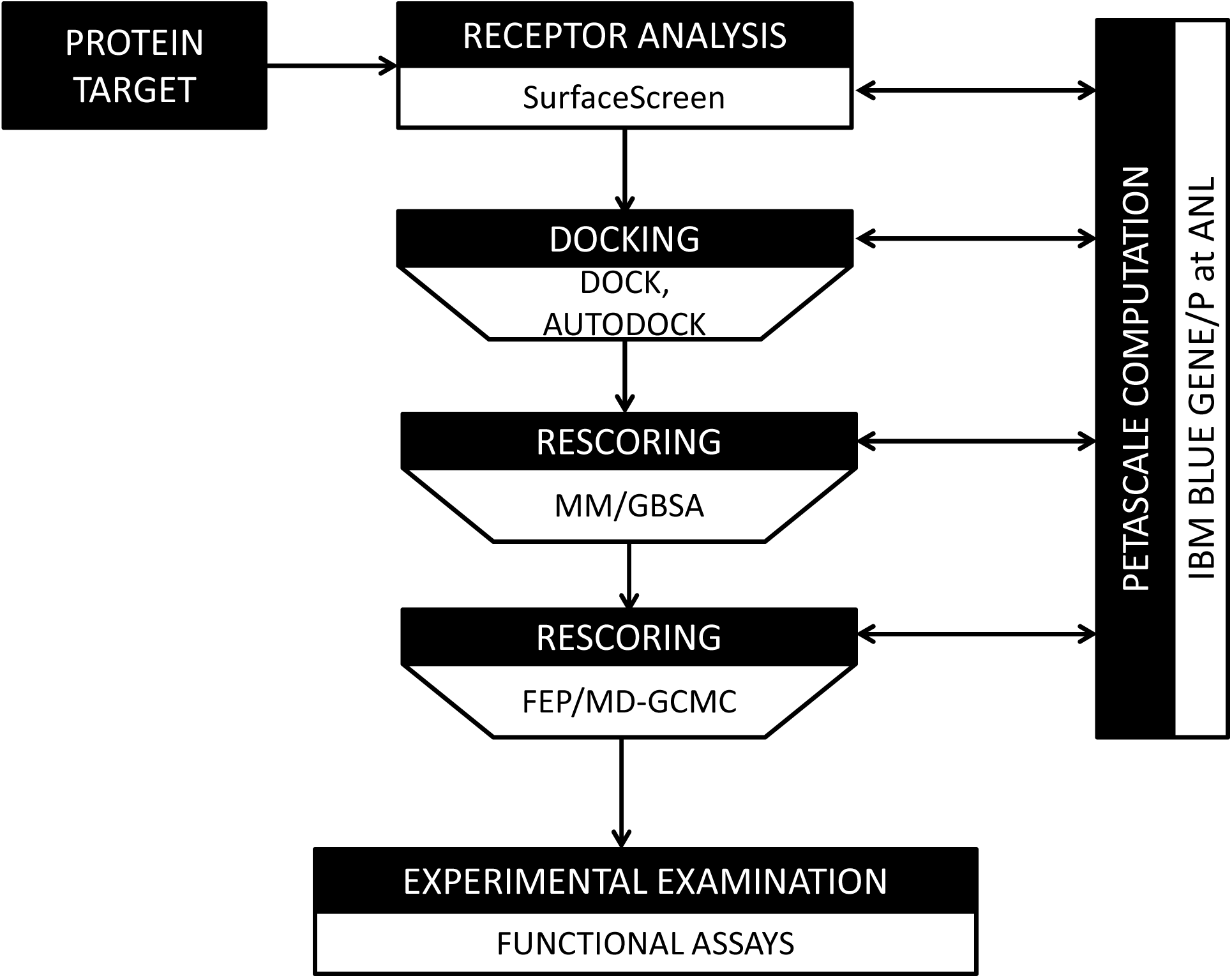
Computational Pipeline Schema for Evaluating Compound Binding.

From this ranked list, we selected 3-5 compounds for further testing. Compounds were then reviewed by experts in medicinal chemistry (KS) and in cancer experimental therapeutics (RB). Factors favoring selection included: availability of compound for purchase (with purity > 90%), higher docking rank, absence of structural characteristics of concern for a therapeutic, such as liable under acid conditions and/or susceptibility to phase I metabolism, especially that due to cytochrome P450, and chemical diversity across compounds for each site. Factors contributing to de-selection included: apparent close analogs to known anti-cancer agents and the possibility that the compound may interfere with a basic and critical biological process (i.e., would have general systemic toxicity), with close analogs of DNA bases representing not uncommon examples. The resultant 38 acquired compounds are listed (**Fig 4**), with 5 each directed against Dxr1, Dxr2, FolC1, FolC2, PRT, TYLT and FabG and 3 against EPSP.

**Fig 4.**
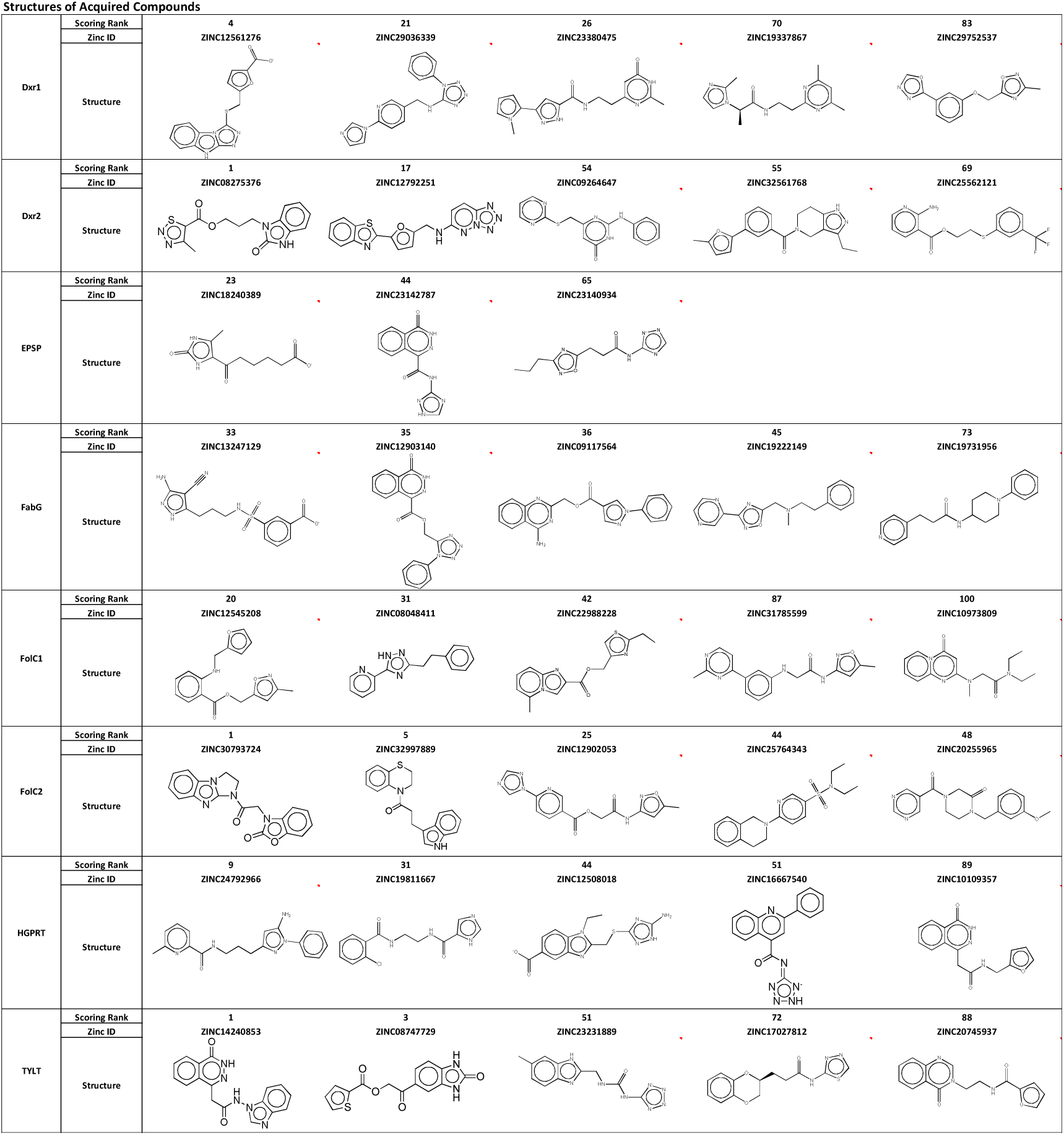
Structure of Acquired Compounds.

### Screening for therapeutic efficacy

We sought to identify compounds that would inhibit the growth of cancer cells, but that also did so selectively. Inhibition of cancer cell growth is a highly sought after attribute of anticancer agents. This was assessed by measuring the ability of a compound to inhibit cell growth in eight-day colony formation assays, examining concentrations ranging from 0.017 to 40 µM. Breast, colon, lung and prostate cancer are four of the most common causes of cancer in the United States, are major causes of cancer cell death, and were examined.[64] Two cell lines for each cancer type were assessed: A549 (adenocarcinoma) and H226 (squamous carcinoma) for lung, MCF-7 (estrogen receptor [ER] positive) and MDA-MB-231 (ER negative) for breast, LNCaP (androgen receptor [AR] positive) and PC3-M (AR negative) for prostate, and HCT116 (wild type p53, mutated KRAS) and HT29 (p53 mutant) for colon.[65]

In parallel with measuring growth inhibition of cancer cell growth, we measured effect on human hematopoietic stem cells. Hematopoietic stem cells reside in the bone marrow and are the source of mature cells present in blood.[66] Bone marrow is one of the most sensitive organs in the body to drug toxicity, especially so for anticancer drugs where it can approach 100%.[67–69] Bone marrow toxicity was measured by performing hematopoietic colony formation assays on human CD34+ stem cells. The stem cell nature of this assay permitted assessment of trilineage colony formation: granulocytic (white blood cells), erythrocytic (red blood cells) and megakaryocytic (platelets), measured by the formation of CFU-GM (colony-forming-granulocyte-macrophage), BFU-E (erythroid burst-forming unites) and CFU-GEMM (colony-forming unit-granulocyte, erythrocyte, monocyte, megakaryocyte) colonies, respectively. Two measurements were taken. Total colonies at eight days were measured, thereby matching the eight-day time frame of cancer cell colony formation assays. At fourteen days, individual colonies have differentiated, permitting quantification of each lineage.

Data for all cancer cell and hematopoietic colony formation assays, across all compound concentrations tested, are provided (**S3 Fig**). We first considered activity at 40 µM, the highest concentration tested. The effects on normal bone marrow, as measured by eight-day stem cell colony formation assay, demonstrated that only 6 (16%) compounds inhibited colony formation by more than 10% and none more than 30% (**Table 1**). Bone marrow colony formation assays provide a measure of toxicity to stem cells and can be extended to fourteen days. If a compound was toxic to stem cells, expanded differences would be expected in the fourteen-day assay as compared to the eight-day. Interestingly, of 6 compounds exerting >10% inhibitory effects at eight days, at fourteen days recovery was observed with 4 compounds and partial recovery with one compound. The overall minimal effects of these compounds on bone marrow function is highlighted by considering the contrasting effect of 5-fluorouracil (5FU). 5FU is an antimetabolite anticancer agent in widespread clinical use and is considered to have only moderate bone marrow suppressive effects clinically.[70] However, 5FU completely suppresses colony formation at 40 µM, and still exerts profound suppressive effects even at concentrations seven fold lower, i.e. 5.7 µM ((**Fig 5**).

**Fig 5.**
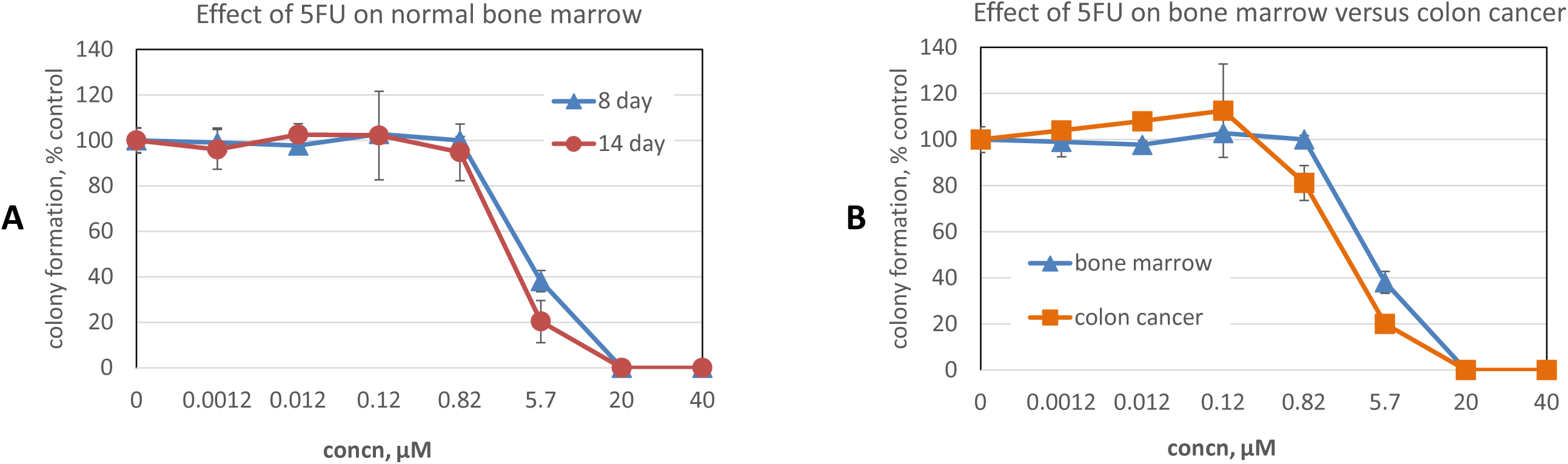
Effect of 5FU on Normal Bone Marrow. (A) Effect of 5FU on eight- and fourteen-day human stem cell hematopoietic colony formation. (B) Effect of 5FU on 8-day colony formation for human stem cells and HT29 colon cancer cells. Data are the mean ± SEM of N=4 and N=2 replicates for stem and HT29 cells, respectively.

**Table 1.** Effect of compounds on normal bone marrow.

|  |  | mean number of colonies, % of control* |  |  |  |  |  |  |  |  |
| --- | --- | --- | --- | --- | --- | --- | --- | --- | --- | --- |
| compound | Dxr2-001 | Dxr2-017 | Dxr2-054 | Dxr2-055 | Dxr2-069 | FolC2-001 | FolC2-005 | FolC2-025 | FolC2-044 | FolC2-048 |
| 8 day | 109 | 70 | 79 | 99 | 108 | 108 | 105 | 93 | 96 | 87 |
| 14 day | 93 | 57 | 112 | 115 | 75 | 87 | 101 | 109 | 73 | 120 |
| compound | FolC1-020 | FolC1-031 | FolC1-042 | FolC1-087 | FolC1-100 | Dxr1-005 | Dxr1-021 | Dxr1-026 | Dxr1-070 | Dxr1-083 |
| 8 day | 107 | 80 | 105 | 107 | 98 | 106 | 101 | 100 | 96 | 97 |
| 14 day | 97 | 85 | 102 | 94 | 96 | 90 | 80 | 128 | 97 | 93 |
| compound | TYLT-001 | TYLT-003 | TYLT-051 | TYLT-072 | TYLT-088 | FabG-033 | FabG-035 | FabG-036 | FabG-045 | FabG-073 |
| 8 day | 117 | 97 | 84 | 88 | 101 | 102 | 113 | 106 | 97 | 97 |
| 14 day | 103 | 94 | 103 | 104 | 112 | 94 | 105 | 100 | 103 | 92 |
| compound | PRT-009 | PRT-031 | PRT-044 | PRT-051 | PRT-P89 | EPSP-023 | EPSP-044 | EPSP-065 |  |  |
| 8 day | 111 | 106 | 101 | 93 | 111 | 94 | 98 | 99 |  |  |
| 14 day | 101 | 97 | 93 | 119 | 93 | 114 | 98 | 98 |  |  |

\*values are mean (N=2 replicates); blue font denotes >10% decrease compared to control

In considering effect on cancer cells, for each compound and each cell line tested we calculated the efficacy ratio, i.e., efficacy ratio = (percent of remaining cancer cell colonies)/(percent of remaining bone marrow colonies), at 8 days after treatment with 40 µM compound (**S4 Fig**). We considered those with an efficacy ratio below 0.5 as active and at least partially selective. The stringency of this is highlighted by considering the effect of 5FU on colon cancer, where it is frequently used clinically in proven curative settings, as well as proven life prolonging settings.[70, 71] At a concentration where 5FU has no effect on bone marrow, i.e. 0.82 µM, the efficacy ratio for human colon cancer HT29 cells is 0.82 (**Fig 5B**). Even at 5.7 µM, where 5FU is toxic to bone marrow, with hematopoietic colony formation at 38% of control, the efficacy ratio for HT29 cells has only dropped to 0.52. In **Table 2** we list all compounds exhibiting an efficacy ratio below 0.5 in at least one cell line. It can be seen that compounds segregate to protein sites. Specifically, of 10 compounds meeting this activity criteria (i.e., efficacy ratio <0.5), 3 were directed at the Dxr2 site, 3 at FolC2, 2 at FolC1, 1 at Dxr1 and 1 at TYLT.

**Table 2.**
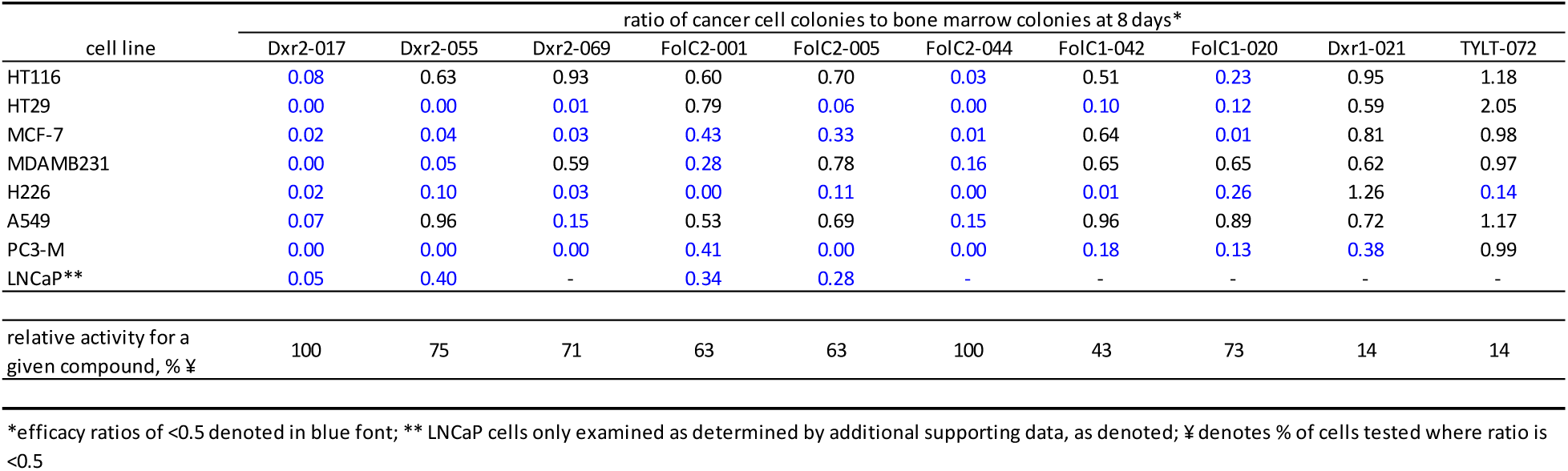
Compounds with desirable efficacy ratios at 40 micromolar.

| cell line | ratio of cancer cell colonies to bone marrow colonies at 8 days* |  |  |  |  |  |  |  |  |  |
| --- | --- | --- | --- | --- | --- | --- | --- | --- | --- | --- |
|  | Dxr2-017 | Dxr2-055 | Dxr2-069 | FolC2-001 | FolC2-005 | FolC2-044 | FolC1-042 | FolC1-020 | Dxr1-021 | TYLT-072 |
| HT116 | 0.08 | 0.63 | 0.93 | 0.60 | 0.70 | 0.03 | 0.51 | 0.23 | 0.95 | 1.18 |
| HT29 | 0.00 | 0.00 | 0.01 | 0.79 | 0.06 | 0.00 | 0.10 | 0.12 | 0.59 | 2.05 |
| MCF-7 | 0.02 | 0.04 | 0.03 | 0.43 | 0.33 | 0.01 | 0.64 | 0.01 | 0.81 | 0.98 |
| MDAMB231 | 0.00 | 0.05 | 0.59 | 0.28 | 0.78 | 0.16 | 0.65 | 0.65 | 0.62 | 0.97 |
| H226 | 0.02 | 0.10 | 0.03 | 0.00 | 0.11 | 0.00 | 0.01 | 0.26 | 1.26 | 0.14 |
| A549 | 0.07 | 0.96 | 0.15 | 0.53 | 0.69 | 0.15 | 0.96 | 0.89 | 0.72 | 1.17 |
| PC3-M | 0.00 | 0.00 | 0.00 | 0.41 | 0.00 | 0.00 | 0.18 | 0.13 | 0.38 | 0.99 |
| LNCaP** | 0.05 | 0.40 | - | 0.34 | 0.28 | - | - | - | - | - |
| relative activity for a given compound, % ¥ | 100 | 75 | 71 | 63 | 63 | 100 | 43 | 73 | 14 | 14 |

\*efficacy ratios of <0.5 denoted in blue font; \*\* LNCaP cells only examined as determined by additional supporting data, as denoted; ¥ denotes % of cells tested where ratio is <0.5

Considered from the perspective of individual protein sites, our findings at 40 µM highlight the importance of Dxr2 and FolC2 sites. As shown in **Table 3** for Dxr2, 60% of tested compounds were considered active against at least one cell line. Of those rated active the efficacy ratio was 0.06 ± 0.022 (mean ± SEM), and for active compounds the percentage of cell lines exhibiting an efficacy ratio below 0.5 ranged from 71-100. For FolC2, 60% of tested compounds were active, the efficacy ratio was 0.15 ± 0.040 (mean ± SEM), and the percentage of susceptible cell lines ranged from 63-100.

**Table 3.** Compound characteristics at each site*.

| site | active compounds, % | activity** |  | range, %‡ |
| --- | --- | --- | --- | --- |
|  |  | mean | SEM |  |
| Dxr2 | 60 | 0.06 | 0.02 | 71-100 |
| FolC2 | 60 | 0.15 | 0.04 | 63-100 |
| FolC1 | 40 | 0.13 | 0.03 | 43-73 |
| Dxr1 | 20 | 0.38 | 0 | 14 |
| TYLT | 20 | 0.14 | 0 | 14 |
| PRT | 0 | NA | NA | NA |
| EPSP | 0 | NA | NA | NA |
| FabG | 0 | NA | NA | NA |

\*compounds at 40 μM
\*\*for compounds with efficacy ratio <0.5, the mean ± SEM ratio is shown
‡for compounds with efficacy ratio <0.5 for a given site, the percentage of cell lines it was active against was calculated, and used to determine range

Examination of compound activity at the next lowest tested concentration of 5.7 µM further supports the importance of Dxr2 directed compounds, and of Dxr2-017 in particular (**Table 4**). Dxr2-017 at 5.7 µM exhibited an efficacy ratio below 0.5 in 75% of cells tested, with those below this cutoff having a 0.25 ± 0.091 (mean ± SEM) efficacy ratio. The activity of Dxr2-017 across all tested concentrations and all cells tested is shown (**Fig 6A, B**), as is its effect on 8- and 14-day bone marrow stem cell trilineage colony formation. At 0.12 µM Dxr2-017 significantly inhibited HT29 colon cancer and MDA-MB-231 breast cancer colony formation by 91.1 ± 4.8% (mean ± SD) and 81.7 ± 15.5%, respectively, compared to control. At concentrations of over 300 fold higher (i.e., 40 µM), Dxr2-017 only inhibited normal bone marrow 8-day colony formation by 30.2 ± 0.7%, and 14-day total colony formation by 42.6 ± 5.8%. The impact on trilineage colony formation can be measured at fourteen days. It demonstrates that 40 µM Dxr2-017 increased CFU-GEMM by 16.1 ± 12.0% compared to control, while CFU-G/M and BFU-E both decreased by 49.8 ± 0.0% and 97.1 ± 4.1%, respectively.

**Fig 6.**
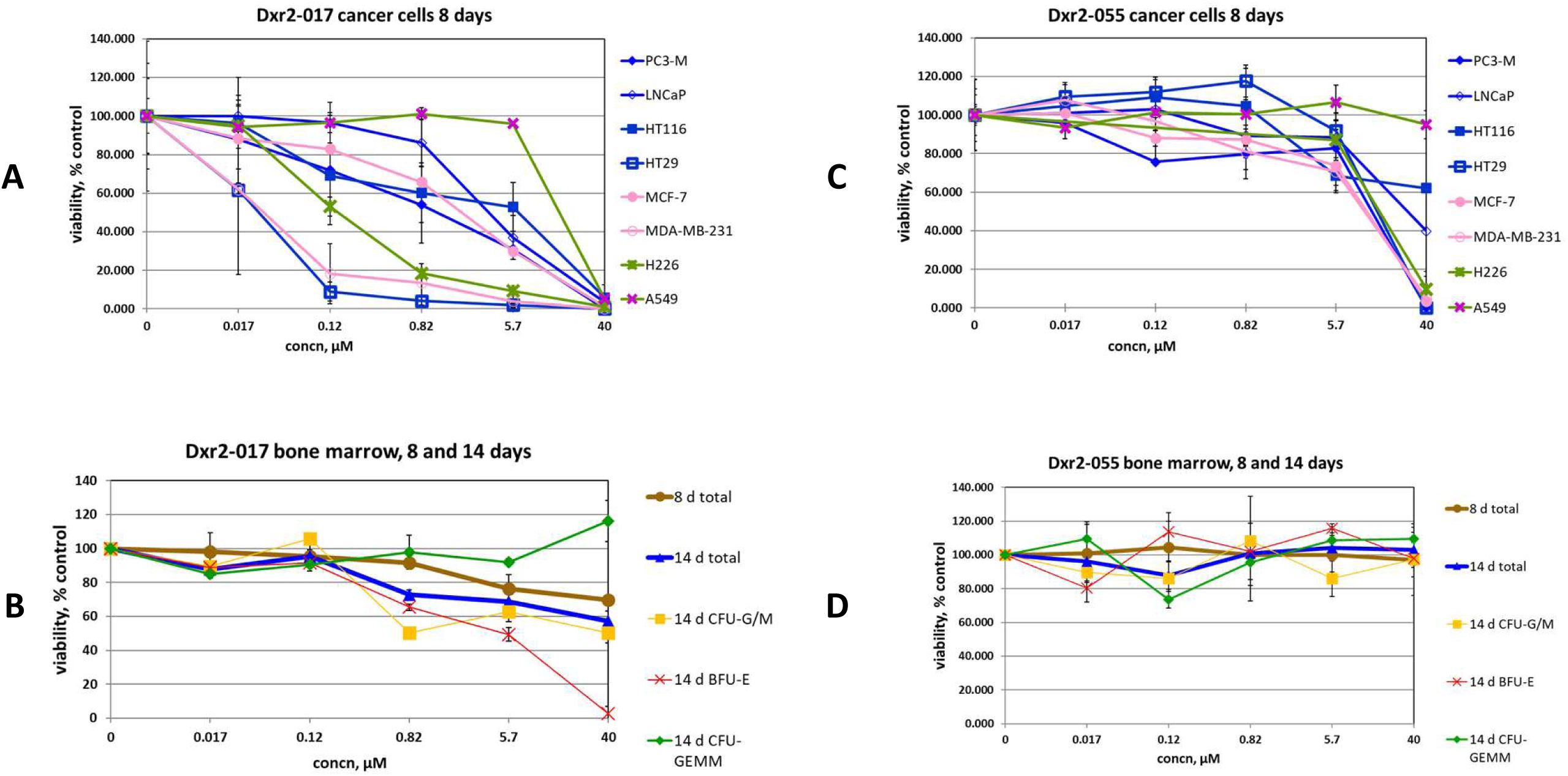
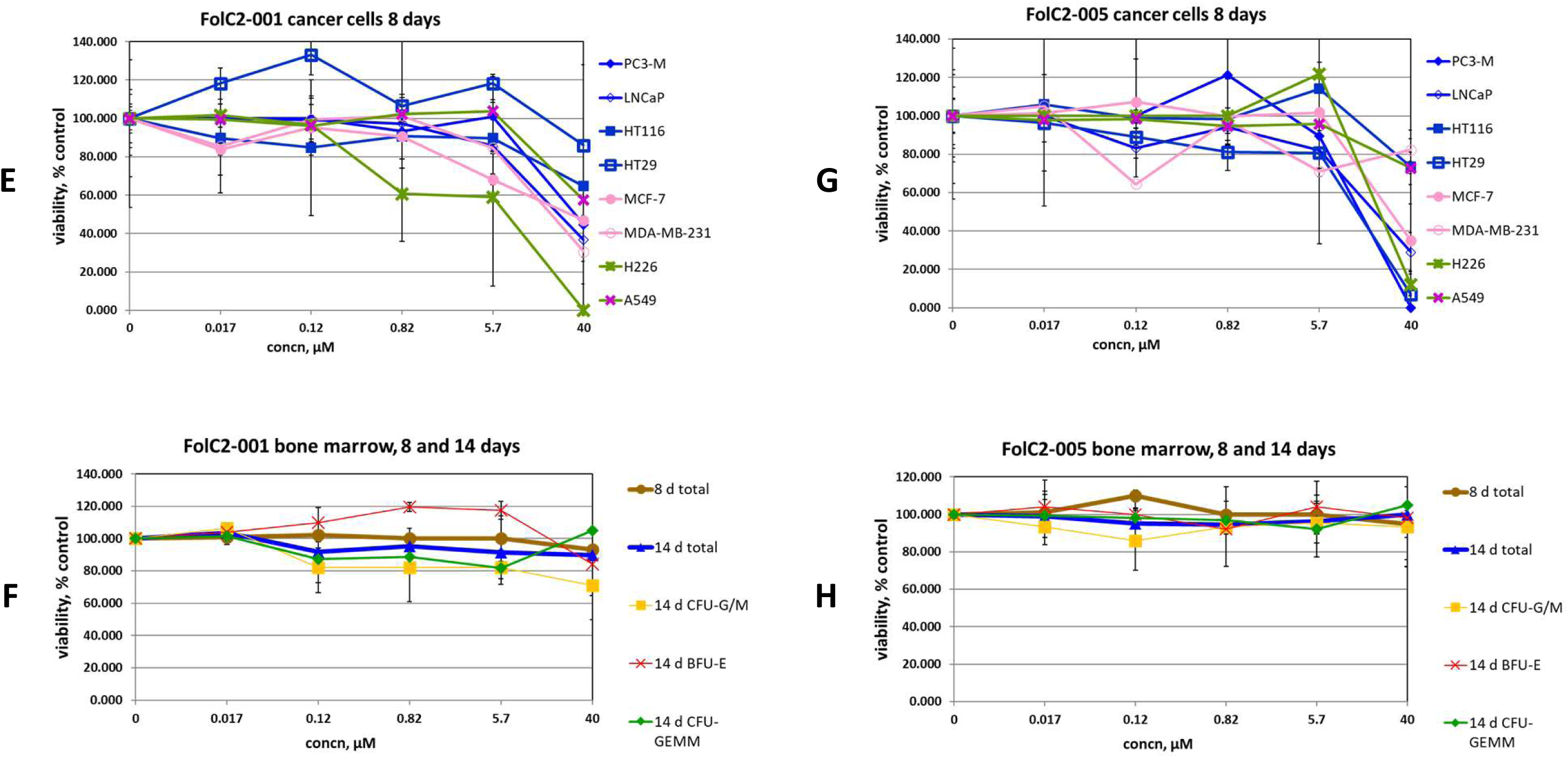
Effect of compounds on cancer cell and bone marrow colony formation. The effect of denoted compounds on eight-day cancer cell colony formation (A, C, E, G). The effect of compounds on eight- and fourteen-day bone marrow colony formation (B, D, F, G). Data are mean ± SD (N=2 replicates), with similar findings in separate experiments (also N=2 replicates).

**Table 4.** Efficacy ratios at 5.7 µM of compound.

| cell line | ratio of cancer cell colonies to bone marrow colonies at 8 days* |  |  |  |
| --- | --- | --- | --- | --- |
|  | Dxr2-017 | FolC2-001 | FolC1-020 | TYLT-072 |
| HT116 | 0.69 | 0.85 | 0.88 | 0.83 |
| HT29 | 0.03 | 1.13 | 0.69 | 1.22 |
| MCF-7 | 0.39 | 0.65 | 0.08 | 0.94 |
| MDAMB231 | 0.05 | 0.81 | 0.96 | 0.95 |
| H226 | 0.12 | 0.56 | 0.96 | 0.26 |
| A549 | 1.26 | 0.99 | 0.84 | 0.99 |
| PC3-M | 0.41 | 0.96 | 0.76 | 1.02 |
| LNCaP** | 0.49 | 0.82 | - | - |
| relative activity for a given compound, % ‡ | 75 | 13 | 14 | 14 |

\* ratios of $<0.5$ denoted in blue font; \*\* LNCaP cells only examined as determined by additional supporting data, as denoted; ‡ denotes % of cells tested where ratio is $<0.5$

Depicted in **Figs 6C-H** are comprehensive data for three additional illustrative compounds, while that for all compounds is shown in **S3 Fig**. Compound Dxr2-055 (**Figs 6C, D**) is directed towards the Dxr2 site and like Dxr2-017 at 40 µM it demonstrates an efficacy ratio below 0.5 against several cancer cells, i.e., 6 cell lines, and has no inhibitory effect on normal bone marrow. However, unlike Dxr2-017, below 40 µM Dxr2-055 efficacy is completely lost. A similar phenomenon is observed with FolC2-005 (**Figs 6G, H**). With FolC2-001 (**Figs 6E, F**), it has the advantage of demonstrating a more dynamic dose-response profile, with 5 cell lines exhibiting an efficacy ratio below 0.5 at 40 µM and some degree of efficacy observed at concentrations below 40 µM. However, FolC2-001’s overall activity is not considered high. At 40 µM for cells where the efficacy ratio is below 0.5, FolC2-001 has an efficacy ratio of 0.29 ± 0.09 (mean ± SEM) which is 9.7 fold higher than that of Dxr2-017, which is 0.03 ± 0.01 (P = 0.001). Further, the activity of FolC2-001 rapidly drops off below 40 µM. The profiles of other compounds (**S3 Fig**) are of even lower interest, mostly based on low efficacy ratios. It is important to consider that this general finding in fact provides additional supportive data related to the selective efficacy of Dxr2-017.

### Expansion of efficacy studies

Above findings support the importance of Dxr2 and FolC2 sites, and of Dxr2-017 and FolC2-001 compounds, respectively. The predicted poses of Dxr2-017 bound to the Dxr2 site and FolC2-001 bound to the FolC2 site are shown (**Fig 7**). The NCI-60 cell line screen evaluates compound growth inhibition and cell toxicity in a highly standardized manner across 60 different cell lines. Dxr2-017 and FolC2-001 were evaluated in this assay (**S5 Fig**). NCI-60 screens evaluate compounds at 10 µM X 48 hours. Findings for both Dxr2-017 and FolC2-001 corroborate those in **Fig 6** indicating Dxr2-017 activity against HT29, MDA-MB-231 and H226, and FolC2-001 against H226. Of high interest, in the NCI-60 screen Dxr2-017 had relatively high activity against melanoma cell lines, inducing cell death in 5 cell lines ranging from 25-65%, and inhibiting growth in 4 cell lines ranging from 76-96%. Its impact on cell death represents the most striking relative difference in response across cell lines.

**Fig 7.**
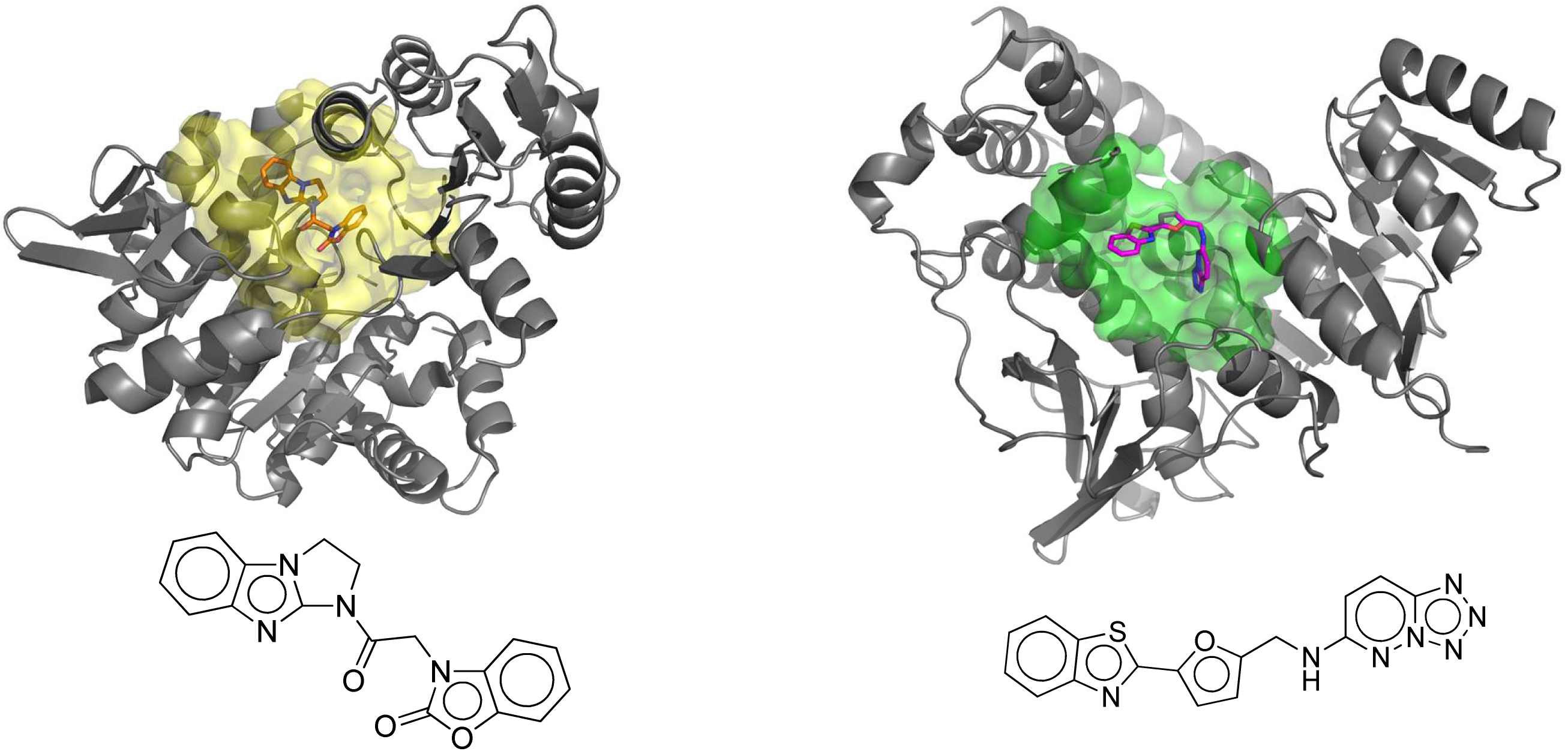
The predicted poses of bound Dxr2-017 and FolC2-001. The poses of FolC2-001 and DxR2-017 bound to their respective FolC2 (yellow) and Dxr2 (green) binding surfaces are depicted, as are ribbon structures of proximal portions of the protein.

We next evaluated the effect of Dxr2-017 on human melanoma M14 and SK-MEL-5 cell lines in eight-day assays (**Fig 8A**). The IC50 values for Dxr2-017 were 19.7 ± 7.4 and 51.7 ± 16.4 (mean ± SEM) nM for M14 and SK-MEL-5, respectively. Compared to eight-day treatment of bone marrow at 40 µM, where colony formation was only decreased by 30%, this translates to greater than 2100-fold efficacy for M14 and 750-fold for SK-MEL-5 cells.

**Fig 8.**
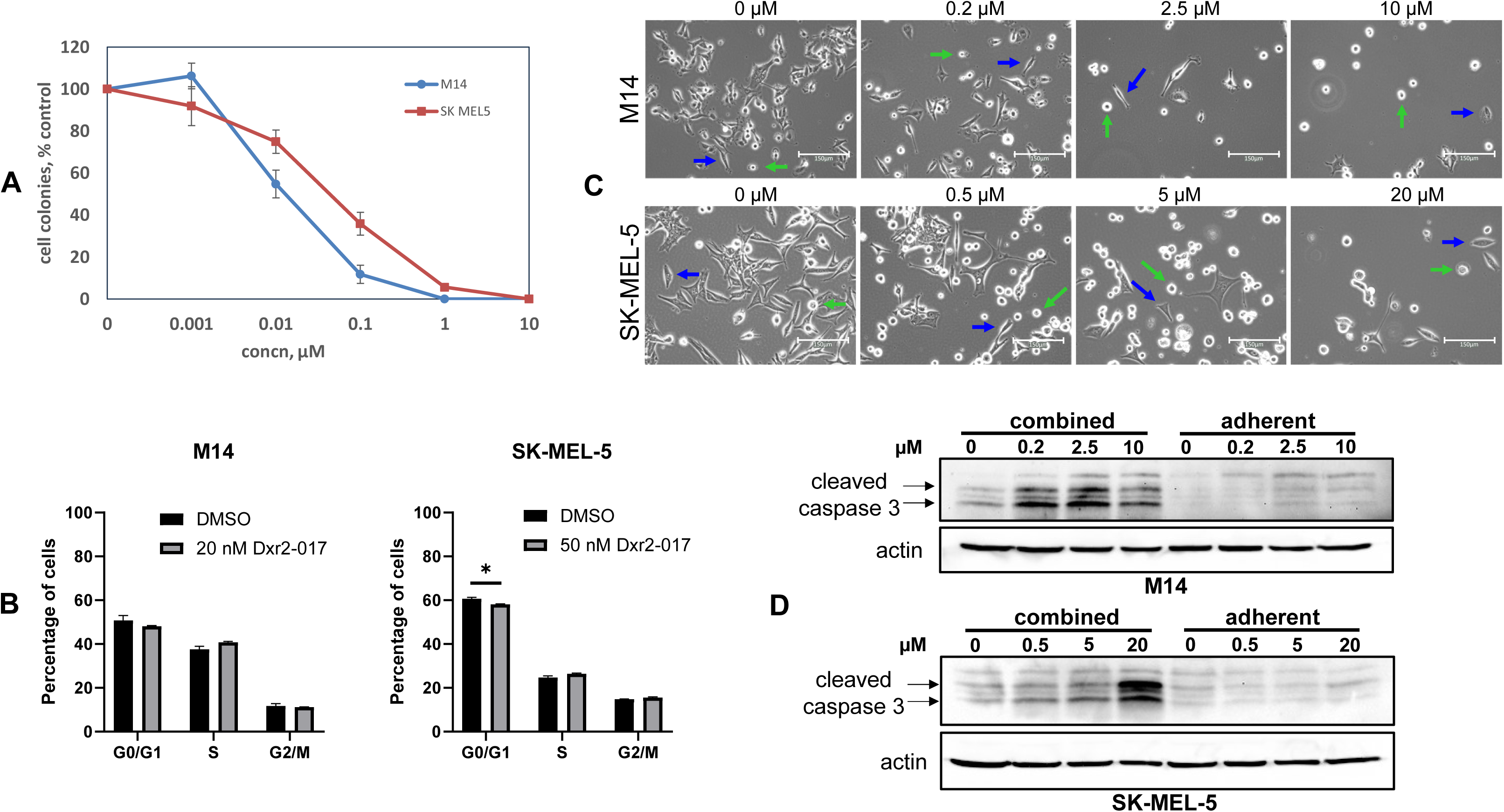
Dxr2-017 inhibits melanoma cell growth through induction of anoikis. (A) Inhibition of melanoma cell growth. Cells were treated with different concentrations of Dxr2-017 and formation of colonies at eight days depicted. Data are expressed as percent of untreated control and are the mean ± SEM (N=3 replicates) of a single experiment. (B) No effect on cycle progression. M14 and SK-MEL-5 cells were treated for 8 days with 20 and 50 nM Dxr2-017, respectively, and phase of cell cycle determined by flow cytometry. Data are the mean ± SEM (N=3). *denotes P-value ≤0.05. (C) Transition to floating cells. Cells were treated for three days with different Dxr2-017 concentrations. Depicted are representative light photomicrographs at 20X; scale bar is150 µm. Blue and green arrows denote adherent and floating cells, respectively. (D) Cleaved caspase 3 is induced in floating cells. Cells were treated for 3 days, cell lysate from only adherent cells, or from adherent and floating cells combined, was probed by Western blot for cleaved caspase 3. All experiments were repeated at separate times, yielding similar results.

We next investigated the manner by which Dxr2-017 inhibited cell growth. We first examined whether Dxr2-017 inhibited cell cycle progression (**Fig 8B**). Treatment of M14 and SK-MEL-5 cells with Dxr2-017 for eight days at concentrations closely matching their respective IC50 values demonstrates little-to-no change in cell cycle parameters.

We then assessed whether Dxr2-017 was inducing apoptosis. Here, we treated M14 and SK-MEL-5 cells with Dxr2-017 for 3 days at concentrations of up 10 and 20 µM, respectively, measured cleaved caspase 3 by Western blot, did so in adherent cells, and did not detect any induction of cleaved caspase 3 (**Fig 8D**). These findings raised the possibility that Dxr2-017 was inducing anoikis, a process in which programmed cell death occurs upon cell detachment.[72, 73] This was supported by demonstrating that with Dxr2-017 treatment there were no morphological differences in adherent cells between treatment and control, but with treatment there were overall less cells with a greater proportion of them becoming floating cells (**Fig 8C, S6 Fig**). As can be seen, these changes were both time- and concentration-dependent. These findings led us to examine cleaved caspase 3 only in adherent cells, as well as in floating and adherent cells combined (**Fig 8D**). While there was no formation of cleaved caspase 3 with up to 10 µM or 20 µM Dxr2-017 in M14 or SK-MEL-5 cells, respectively, when only the adherent fraction was probed, in contrast clear formation was observed when the floating cell fraction was included.

## Discussion

We demonstrate proof-of-concept that 3D protein structure represents an important source of primary information that can be used to begin the process of drug discovery. We elected to use a protein structure library created by us, as well as a suite of analytic tools created by us. Finally, we elected to apply this strategy to the task of discovering anticancer agents that selectively inhibited cancer cell growth. The basic concept of recognizing that novel protein structure represents an information portal to novel function represents a rich strategy that is broadly adaptable. In particular, larger libraries are available, are increasingly accessible, exponential advances in protein computational analytics continue to emerge, and applications range across the spectrum of the most basic biology to an array of given end-use functions.[74–79]

Our demonstration that compounds selected as active in the current study segregated to specific protein pockets is of high importance. Recognizing that while our current strategy used physically defined protein structure and this was coupled to experimental measures of function, the intermediate process of compound screening was virtual. That our functional measures of activity segregated to protein pockets provides an important measure of the real-world utility of our computational approaches as applied to virtual screening.

Further, we consider our integrated and layered screening approach to be of high value. The first layer in this approach used up front sets of computational analytics. The second layer transitioned to experimental analytics and involved in-parallel selection and de-selection screens. The latter has been a strategy previously employed by us, which was also highly successful.[65] At this point our analytics yielded priority compounds, with Dxr2-017 emerging as a clear lead. Our incorporation at this stage of a broader screening layer, i.e., the NCI-60 cell line screen, further solidified the lead and served to redirect our disease focus to melanoma.

Our demonstration that Dxr2-017 was potent, specific and acting through a novel mechanism represent three individual factors that independently, and certainly together, support our hypothesis that unique protein structure provides a high value entry point for the downstream discovery of novel therapeutics. Specifically, IC50 values in the low nM range highlight the potency of Dxr2-017. That concentrations over 2100-fold higher have little effect on bone marrow stem cells provides a powerful measure of specificity. While comprehensive examination of all systemic organ systems over an extended time is necessary to provide a more comprehensive assessment of specificity, given the almost universally high sensitivity of bone marrow to anticancer agents, and the magnitude of an over 2100-fold therapeutic window, high specificity is well supported. The novelty of Dxr2-017 is supported by its induction of anoikis. Although anoikis has long been recognized as an important cellular process, there is a limited understanding of its regulation, and a great need for therapeutics that specifically induce it.[72, 73, 80, 81] Findings from the current study provide a new means to specifically target anoikis. Further, the small chemical nature of Dxr2-017 also represents a new tool that can be used as a probe to interrogate biology. It is recognized that large knowledge gaps exist with respect to further advancing Dxr2-017 along a drug development pathway. A major one relates to its actual target(s). In this regard it should be noted that the Dxr2 pocket is part the protein 1-deoxy-D-xylulose 5-phosphate reductoisomerase, also known as DXP reductoisomerase, and represents a protein not expressed in humans.

There are several limitations in the current study. Our selection methodology of proteins to be analyzed relied heavily on human-based selection factors. Our virtual analytics did not employ newer artificial intelligence-based techniques. While such cutting edge techniques have the clear potential to add, they do need to be viewed with caution.[74, 82] The number of chemicals subjected to experimental analysis was small. While this latter factor could also be considered a measure of efficiency, high throughput methods would likely serve as additive. While our approach of considering phylogenetic preservation of structure has a justifiable rational basis, it is not possible within the current study to make any strong attributive statements as to its overall importance. Our findings could be considered as providing additional evidence to support the hypothesis that such considerations are important, at least as they relate to identifying sought after therapeutic effects. Current findings do support future investigations probing this notion.

This study provides proof-of-concept that 3D protein structure represents an important starting point from which to launch the drug discovery process. The basic nature of this strategy has wider implications related to application and biology. Dxr2-017 was identified as a highly active compound that inhibits the growth of human melanoma cells, has a minimal impact on human bone marrow, induces anoikis, and holds high promise as a therapeutic, or for informing the development of such.

## Materials and Methods

### Chemicals and cell lines

Compounds were purchased from manufacturers (**S7 Fig**). The following human cells were obtained from American Type Culture Collection: breast cancer, MCF-7, MDA-MB-231; lung cancer, H226, A549; colon cancer, HT116, HT29; prostate cancer, LNCaP; melanoma, M14, SK-MEL-5. The origin and characteristics of PC3-M cells were previously described by us.[83] Cells were cultured as previously described by us.[84, 85] Briefly, cells were drawn from frozen stocks, periodically replenished on a standard basis, grown in 5% CO_2_, 37°C, maintained under sub confluent exponential growth conditions, passaged 1-2 times weekly, manipulated one cell line at a time, with complete sterilization of working surfaces in between, and routinely monitored for Mycoplasma (PlasmoTest™, InvivoGen, San Diego, CA). Authentication procedures involved: after acquisition from the originator, grown and expanded under quarantine conditions, stored as primary stocks and not used until confirmed mycoplasma negative; morphologic examination confirmed expected phenotype; growth characteristics confirmed; hormone responsiveness or lack thereof confirmed.

### Development and characterization of a protein x-ray crystallographic library

Using the Argonne National Laboratory Advanced Proton Source, we have generated and characterized a high resolution x-ray crystallographic protein library as described by us.[13] All final structures have been deposited in the Protein Data Bank (PDB).[86]

### Development and validation of a physics-based protein structure analysis platform

We have developed a physics-based protein structure analysis platform, allowing consideration of dynamic protein flexibility and the influence of solvent, that was scaled to run on a supercomputer, thus providing the capability to probe large protein libraries for novel structural motifs. Specifically, we integrated a suite of methodologies, i.e., CASTp, pvSOAR, SurfaceAlign, SurfaceScreen, scaled to operate on the 163,840 core Intrepid BlueGene/P (Intrepid BG/P) supercomputer, of the Argonne National Laboratory Advanced Leadership Computing Facility (ALCF), and organized them into a computational pipeline designed to identify, characterize and compare protein surfaces.[14–20]

### In silico compound screening

We have constructed a computational pipeline for docking and scoring compound interactions with protein pockets.[15–17, 60–63] Using SurfaceScreen methodology, our pipeline initially incorporated AutoDock[60] and DOCK[61] docking applications, together providing a broad coverage of protein-small molecule docking strategies, including accounting for protein residue and compound flexibility. Top ranking initial docking poses are then ‘funneled’ into increasingly more sophisticated and computationally complex approximations of free energy of interaction using the MM-GBSA[62] (molecular mechanics-generalized Born surface area) and FEP/MD-GCMC[63] (molecular dynamics free energy perturbation-grand canonical Monte Carlo) methods, with rescoring of poses conducted in a hierarchical fashion.

### Growth Assays

Eight-day colony formation assays of cancer cells were performed as described by us.[87] Briefly, 0.5-1.0 x 10^3^ cells were seeded into a 12 well plate, after 24 hours compound added and eight days later colonies (of 3 or more cells) were scored after trypan blue staining. Conditions were in replicates of N = 2. All compounds were stored under desiccated conditions in a sealed container at minus 20°C upon arrival. One day prior to use, they were placed into DMSO, stored at 20°C as stock solutions, thawed just prior to use, and not reused.

### Bone marrow stem cell colony formation assays

Human hematopoietic stem cell trilineage colony formation assays were performed as previously described by us.[65] Briefly, human cord blood CD34+ cells (catalog #: 70008.5) were purchased from StemCell Technology Inc., and seeded at 1 x 10^4^ cells per 35 mm dish. Cells were cultured in MethoCult™ Express (StemCell Technologies, Inc.) semisolid media, per manufacturer’s instructions, in the presence of compound or vehicle control. At eight days colonies, i.e., groups of ≥ 10 cells, were counted. At fourteen days, individual colonies of BFU-E, CFU-GM and CFU-GEMM were identified based on manufacturers’ morphologic characterization criteria. Assays were run in replicates of N = 2; some were run in replicates of N = 3, as denoted.

### Cell cycle analysis

A total of 1 x 10^4^ M14 cells and 1 x 10^4^ SK-MEL-5 cells were seeded into 10 cm culture dishes. Twenty-four hours later cells, in replicates of N = 3, were treated with the Dxr2-017 at 20 nM for M14 and 50 nM for SK-MEL-5, or vehicle control for a total of eight days, with replacement of media, along with <u>Dxr2-017 or vehicle</u>, at day five. Cells were then collected, filtered to obtain single cell suspensions, equal numbers of cells fixed in 1 ml 70% ethanol at 4°C for 1 hour, washed with PBS, and resuspended in 1 mL Telford reagent (1 mM EDTA disodium salt, 2.5 U/ml RNAse A, 75 µM Propidium Iodide, 0.1% Triton X-100 in PBS), incubated at 4°C overnight. Cells were analyzed by flow cytometry (FACSCalibur3 analyzer, BD Biosciences), and the percentage of cells within G0/G1, S, and G2/M phases were determined via analysis using Modfit LT 4.0.5.

### Western blot and image acquisition

Briefly, M14 and SK-MEL-5 cells were treated with different concentrations of Dxr2-017 or vehicle, for 3 or 8 days, as denoted. Before sample collection, cell images were captured at 20X in transmitted light channel using EVOS™ M5000 Imaging System (Invitrogen). Cells were collected and lysed using RIPA lysis buffer (Thermo Scientific^TM^, #89900) (25 mM Tris-HCl pH 7.6, 150 mM NaCl, 1% NP-40, 1% sodium deoxycholate, 0.1% SDS) supplemented with protease and phosphatase inhibitor cocktail (Thermo Scientific^TM^, #78442) and 1 mM PMSF, and Western blot performed on resultant protein as described by us.[85] Samples were probed using antibody against cleaved caspase 3 (Cell Signaling Technology, #9664) and β-actin (Santa Cruz, #sc-47778).

## Acknowledgements

Financial support for this research was provided to R.B. by the Gerald G. Pabst Trust fund, to A.B. from the National Institutes of Health (GM094585), resources of the Argonne Leadership Computing Facility at Argonne National Laboratory (supported by the Office of Science of the US Department of Energy under contract DE-AC02-06CH11357) and allocations for computing, supported by the Department of Energy’s Innovative and Novel Computational Impact on Theory and Experiment (INCITE) program. The Center for Structural Genomics of Infectious Diseases has been funded in whole or in part with federal funds from the National Institute of Allergy and Infectious Diseases, National Institutes of Health, Department of Health and Human Services, under Contract Nos. HHSN272200700058C and HHSN272201200026C (to W.F.A). The authors thank Jaclyn Hollinger for technical support, and the University of Nebraska Medical Center Flow Cytometry core facility for project support.

## References

1. Zhung W, Kim H, Kim WY. 3D molecular generative framework for interaction-guided drug design. Nat Commun. 2024;15(1):2688. Epub 20240327. doi: 10.1038/s41467-024-47011-2 PubMed PMID: 38538598; PubMed Central PMCID: PMCPMC10973397.

2. Sakuma K, Kobayashi N, Sugiki T, Nagashima T, Fujiwara T, Suzuki K, et al. Design of complicated all-alpha protein structures. Nat Struct Mol Biol. 2024;31(2):275–82. Epub 20240104. doi: 10.1038/s41594-023-01147-9. PubMed PMID: 38177681.

3. Seoane B, Carbone A. The complexity of protein interactions unravelled from structural disorder. PLoS Comput Biol. 2021;17(1):e1008546. Epub 20210108. doi: 10.1371/journal.pcbi.1008546. PubMed PMID: 33417598; PubMed Central PMCID: PMCPMC7846008.

4. Yao B, Li Z, Ding M, Chen M. Three-dimensional protein model similarity analysis based on salient shape index. BMC Bioinformatics. 2016;17:131. Epub 20160318. doi: 10.1186/s12859-016-0983-z. PubMed PMID: 26987968; PubMed Central PMCID: PMCPMC4797110.

5. Batool M, Ahmad B, Choi S. A Structure-Based Drug Discovery Paradigm. Int J Mol Sci. 2019;20(11). Epub 20190606. doi: 10.3390/ijms20112783. PubMed PMID: 31174387; PubMed Central PMCID: PMCPMC6601033.

6. Mohs RC, Greig NH. Drug discovery and development: Role of basic biological research. Alzheimers Dement (N Y). 2017;3(4):651–7. Epub 20171111. doi: 10.1016/j.trci.2017.10.005. PubMed PMID: 29255791; PubMed Central PMCID: PMCPMC5725284.

7. Barnash KD, James LI, Frye SV. Target class drug discovery. Nat Chem Biol. 2017;13(10):1053–6. doi: 10.1038/nchembio.2473. PubMed PMID: 28926557; PubMed Central PMCID: PMCPMC5973815.

8. Childers WE, Elokely KM, Abou-Gharbia M. The Resurrection of Phenotypic Drug Discovery. ACS Med Chem Lett. 2020;11(10):1820–8. Epub 20200306. doi: 10.1021/acsmedchemlett.0c00006. PubMed PMID: 33062159; PubMed Central PMCID: PMCPMC7549108.

9. Hingorani AD, Kuan V, Finan C, Kruger FA, Gaulton A, Chopade S, et al. Improving the odds of drug development success through human genomics: modelling study. Sci Rep. 2019;9(1):18911. Epub 20191211. doi: 10.1038/s41598-019-54849-w. PubMed PMID: 31827124; PubMed Central PMCID: PMCPMC6906499.

10. Sun D, Gao W, Hu H, Zhou S. Why 90% of clinical drug development fails and how to improve it? Acta Pharm Sin B. 2022;12(7):3049–62. Epub 20220211. doi: 10.1016/j.apsb.2022.02.002. PubMed PMID: 35865092; PubMed Central PMCID: PMCPMC9293739.

11. Schlander M, Hernandez-Villafuerte K, Cheng CY, Mestre-Ferrandiz J, Baumann M. How Much Does It Cost to Research and Develop a New Drug? A Systematic Review and Assessment. Pharmacoeconomics. 2021;39(11):1243–69. Epub 20210809. doi: 10.1007/s40273-021-01065-y. PubMed PMID: 34368939; PubMed Central PMCID: PMCPMC8516790.

12. Wouters OJ, McKee M, Luyten J. Estimated Research and Development Investment Needed to Bring a New Medicine to Market, 2009-2018. JAMA. 2020;323(9):844–53. doi: 10.1001/jama.2020.1166. PubMed PMID: 32125404; PubMed Central PMCID: PMCPMC7054832.

13. Stacy R, Anderson WF, Myler PJ. Structural Genomics Support for Infectious Disease Drug Design. ACS Infect Dis. 2015;1(3):127–9. Epub 20150120. doi: 10.1021/id500048p. PubMed PMID: 25984568; PubMed Central PMCID: PMCPMC4426352.

14. Binkowski TA, Joachimiak A. Protein functional surfaces: global shape matching and local spatial alignments of ligand binding sites. BMC Struct Biol. 2008;8:45. Epub 2008/10/29. doi: 1472-6807-8-45 [pii] 10.1186/1472-6807-8-45. PubMed PMID: 18954462; PubMed Central PMCID: PMC2626596.

15. Dundas J, Ouyang Z, Tseng J, Binkowski A, Turpaz Y, Liang J. CASTp: computed atlas of surface topography of proteins with structural and topographical mapping of functionally annotated residues. Nucleic Acids Res. 2006;34(Web Server issue):W116–8. Epub 2006/07/18. doi: 34/suppl_2/W116 [pii] 10.1093/nar/gkl282. PubMed PMID: 16844972; PubMed Central PMCID: PMC1538779.

16. Binkowski TA, Freeman P, Liang J. pvSOAR: detecting similar surface patterns of pocket and void surfaces of amino acid residues on proteins. Nucleic Acids Res. 2004;32(Web Server issue):W555–8. Epub 2004/06/25. doi: 10.1093/nar/gkh39032/suppl_2/W555 [pii]. PubMed PMID: 15215448; PubMed Central PMCID: PMC441528.

17. Binkowski TA, Naghibzadeh S, Liang J. CASTp: Computed Atlas of Surface Topography of proteins. Nucleic Acids Res. 2003;31(13):3352–5. Epub 2003/06/26. PubMed PMID: 12824325; PubMed Central PMCID: PMC168919.

18. Binkowski TA, Cuff M, Nocek B, Chang C, Joachimiak A. Assisted assignment of ligands corresponding to unknown electron density. J Struct Funct Genomics. 2010;11(1):21–30. Epub 2010/01/22. doi: 10.1007/s10969-010-9078-7. PubMed PMID: 20091237.

19. Binkowski TA, Joachimiak A, Liang J. Protein surface analysis for function annotation in high-throughput structural genomics pipeline. Protein Sci. 2005;14(12):2972–81. Epub 2005/12/03. doi: 14/12/2972 [pii] 10.1110/ps.051759005. PubMed PMID: 16322579; PubMed Central PMCID: PMC2253251.

20. Binkowski TA, Adamian L, Liang J. Inferring functional relationships of proteins from local sequence and spatial surface patterns. J Mol Biol. 2003;332(2):505–26. Epub 2003/09/02. doi: S0022283603008829 [pii]. PubMed PMID: 12948498.

21. Binkowski TA, Jiang W, Roux B, Anderson WF, Joachimiak A. Virtual high-throughput ligand screening. Methods Mol Biol. 2014;1140:251–61. Epub 2014/03/05. doi: 10.1007/978-1-4939-0354-2_19. PubMed PMID: 24590723; PubMed Central PMCID: PMCPMC4073479.

22. Dyda F, Klein DC, Hickman AB. GCN5-related N-acetyltransferases: a structural overview. Annu Rev Biophys Biomol Struct. 2000;29:81–103. Epub 2000/08/15. doi: 10.1146/annurev.biophys.29.1.81. PubMed PMID: 10940244; PubMed Central PMCID: PMCPMC4782277.

23. Chakravarti IM, Laha RG, Roy J. Handbook of Methods of Applied Statistics: olume I. John Wiley and Sons; 1967.

24. Umeyama S. Least-squares estimation of transformation paremeters between two point patterns. IEEE Trans Pattern Anal Mach Intell. 1991;13:376–80.

25. Liang J, Edelsbrunner H, Fu P, Sudhakar PV, Subramaniam S. Analytical shape computation of macromolecules: II. Inaccessible cavities in proteins. Proteins. 1998;33(1):18–29. Epub 1998/09/19. PubMed PMID: 9741841.

26. Liang J, Edelsbrunner H, Fu P, Sudhakar PV, Subramaniam S. Analytical shape computation of macromolecules: I. Molecular area and volume through alpha shape. Proteins. 1998;33(1):1–17. Epub 1998/09/19. PubMed PMID: 9741840.

27. Liang J, Edelsbrunner H, Woodward C. Anatomy of protein pockets and cavities: measurement of binding site geometry and implications for ligand design. Protein Sci. 1998;7(9):1884–97. Epub 1998/10/07. doi: 10.1002/pro.5560070905. PubMed PMID: 9761470; PubMed Central PMCID: PMCPMC2144175.

28. Rush TS, 3rd, Grant JA, Mosyak L, Nicholls A. A shape-based 3-D scaffold hopping method and its application to a bacterial protein-protein interaction. J Med Chem. 2005;48(5):1489-95. Epub 2005/03/04. doi: 10.1021/jm040163o. PubMed PMID: 15743191.

29. Feinstein WP, Brylinski M. Calculating an optimal box size for ligand docking and virtual screening against experimental and predicted binding pockets. J Cheminform. 2015;7:18. Epub 2015/06/18. doi: 10.1186/s13321-015-0067-5. PubMed PMID: 26082804; PubMed Central PMCID: PMCPMC4468813.

30. Muegge I. Pharmacophore features of potential drugs. Chemistry. 2002;8(9):1976–81. Epub 2002/05/01. doi: 10.1002/1521-3765(20020503)8:9<1976::AID-CHEM1976>3.0.CO;2-K. PubMed PMID: 11981881.

31. Coleman RG, Sharp KA. Protein pockets: inventory, shape, and comparison. J Chem Inf Model. 2010;50(4):589–603. Epub 2010/03/09. doi: 10.1021/ci900397t. PubMed PMID: 20205445; PubMed Central PMCID: PMCPMC2859996.

32. Bastolla U, Porto M, Eduardo Roman MH, Vendruscolo MH. Connectivity of neutral networks, overdispersion, and structural conservation in protein evolution. Journal of molecular evolution. 2003;56(3):243–54. Epub 2003/03/04. doi: 10.1007/s00239-002-2350-0. PubMed PMID: 12612828.

33. Orengo CA, Bray JE, Buchan DW, Harrison A, Lee D, Pearl FM, et al. The CATH protein family database: a resource for structural and functional annotation of genomes. Proteomics. 2002;2(1):11–21. Epub 2002/01/15. doi: 10.1002/1615-9861(200201)2:1<11::AID-PROT11>3.0.CO;2-T [pii]. PubMed PMID: 11788987.

34. Erdin S, Ward RM, Venner E, Lichtarge O. Evolutionary trace annotation of protein function in the structural proteome. J Mol Biol. 2010;396(5):1451–73. Epub 2009/12/29. doi: S0022-2836(09)01544-7 [pii] 10.1016/j.jmb.2009.12.037. PubMed PMID: 20036248; PubMed Central PMCID: PMC2831211.

35. Madabushi S, Yao H, Marsh M, Kristensen DM, Philippi A, Sowa ME, et al. Structural clusters of evolutionary trace residues are statistically significant and common in proteins. J Mol Biol. 2002;316(1):139–54. Epub 2002/02/07. doi: 10.1006/jmbi.2001.5327 S0022283601953276 [pii]. PubMed PMID: 11829509.

36. Lichtarge O, Bourne HR, Cohen FE. An evolutionary trace method defines binding surfaces common to protein families. J Mol Biol. 1996;257(2):342–58. Epub 1996/03/29. doi: S0022-2836(96)90167-9 [pii] 10.1006/jmbi.1996.0167. PubMed PMID: 8609628.

37. Landau M, Mayrose I, Rosenberg Y, Glaser F, Martz E, Pupko T, et al. ConSurf 2005: the projection of evolutionary conservation scores of residues on protein structures. Nucleic Acids Res. 2005;33(Web Server issue):W299–302. Epub 2005/06/28. doi: 33/suppl_2/W299 [pii] 10.1093/nar/gki370. PubMed PMID: 15980475; PubMed Central PMCID: PMC1160131.

38. La D, Sutch B, Livesay DR. Predicting protein functional sites with phylogenetic motifs. Proteins. 2005;58(2):309–20. Epub 2004/12/02. doi: 10.1002/prot.20321. PubMed PMID: 15573397.

39. del Sol A, Pazos F, Valencia A. Automatic methods for predicting functionally important residues. J Mol Biol. 2003;326(4):1289–302. Epub 2003/02/19. doi: S0022283602014511 [pii]. PubMed PMID: 12589769.

40. Balendiran GK, Molina JA, Xu Y, Torres-Martinez J, Stevens R, Focia PJ, et al. Ternary complex structure of human HGPRTase, PRPP, Mg2+, and the inhibitor HPP reveals the involvement of the flexible loop in substrate binding. Protein Sci. 1999;8(5):1023–31. Epub 1999/05/25. doi: 10.1110/ps.8.5.1023. PubMed PMID: 10338013; PubMed Central PMCID: PMCPMC2144341.

41. Keough DT, Hockova D, Holy A, Naesens LM, Skinner-Adams TS, Jersey J, et al. Inhibition of hypoxanthine-guanine phosphoribosyltransferase by acyclic nucleoside phosphonates: a new class of antimalarial therapeutics. J Med Chem. 2009;52(14):4391–9. Epub 2009/06/17. doi: 10.1021/jm900267n. PubMed PMID: 19527031.

42. Ansari MY, Dikhit MR, Sahoo GC, Das P. Comparative modeling of HGPRT enzyme of L. donovani and binding affinities of different analogs of GMP. Int J Biol Macromol. 2012;50(3):637–49. Epub 2012/02/14. doi: 10.1016/j.ijbiomac.2012.01.010. PubMed PMID: 22327112.

43. Ansari MY, Equbal A, Dikhit MR, Mansuri R, Rana S, Ali V, et al. Establishment of correlation between in-silico and in-vitro test analysis against Leishmania HGPRT to inhibitors. Int J Biol Macromol. 2016;83:78–96. Epub 2015/12/01. doi: 10.1016/j.ijbiomac.2015.11.051. PubMed PMID: 26616453.

44. Bognar AL, Osborne C, Shane B, Singer SC, Ferone R. Folylpoly-gamma-glutamate synthetase-dihydrofolate synthetase. Cloning and high expression of the Escherichia coli folC gene and purification and properties of the gene product. J Biol Chem. 1985;260(9):5625–30. Epub 1985/05/10. PubMed PMID: 2985605.

45. Wang P, Wang Q, Yang Y, Coward JK, Nzila A, Sims PF, et al. Characterisation of the bifunctional dihydrofolate synthase-folylpolyglutamate synthase from Plasmodium falciparum; a potential novel target for antimalarial antifolate inhibition. Mol Biochem Parasitol. 2010;172(1):41–51. Epub 2010/03/31. doi: 10.1016/j.molbiopara.2010.03.012. PubMed PMID: 20350571; PubMed Central PMCID: PMCPMC2877875.

46. Takahashi S, Kuzuyama T, Watanabe H, Seto H. A 1-deoxy-D-xylulose 5-phosphate reductoisomerase catalyzing the formation of 2-C-methyl-D-erythritol 4-phosphate in an alternative nonmevalonate pathway for terpenoid biosynthesis. Proc Natl Acad Sci U S A. 1998;95(17):9879–84. Epub 1998/08/26. doi: 10.1073/pnas.95.17.9879. PubMed PMID: 9707569; PubMed Central PMCID: PMCPMC21430.

47. Deng L, Diao J, Chen P, Pujari V, Yao Y, Cheng G, et al. Inhibition of 1-deoxy-D-xylulose-5-phosphate reductoisomerase by lipophilic phosphonates: SAR, QSAR, and crystallographic studies. J Med Chem. 2011;54(13):4721–34. Epub 2011/05/13. doi: 10.1021/jm200363d. PubMed PMID: 21561155; PubMed Central PMCID: PMCPMC3601441.

48. Liu Y, Feng Y, Cao X, Li X, Xue S. Structure-directed construction of a high-performance version of the enzyme FabG from the photosynthetic microorganism Synechocystis sp. PCC 6803. FEBS Lett. 2015;589(20 Pt B):3052-7. Epub 2015/09/12. doi: 10.1016/j.febslet.2015.09.001. PubMed PMID: 26358291.

49. Hoang TT, Sullivan SA, Cusick JK, Schweizer HP. Beta-ketoacyl acyl carrier protein reductase (FabG) activity of the fatty acid biosynthetic pathway is a determining factor of 3-oxo-homoserine lactone acyl chain lengths. Microbiology (Reading). 2002;148(Pt 12):3849–56. Epub 2002/12/14. doi: 10.1099/00221287-148-12-3849. PubMed PMID: 12480888.

50. Gurvitz A. The essential mycobacterial genes, fabG1 and fabG4, encode 3-oxoacyl-thioester reductases that are functional in yeast mitochondrial fatty acid synthase type 2. Mol Genet Genomics. 2009;282(4):407–16. Epub 2009/08/18. doi: 10.1007/s00438-009-0474-2. PubMed PMID: 19685079; PubMed Central PMCID: PMCPMC2746893.

51. Ducasse-Cabanot S, Cohen-Gonsaud M, Marrakchi H, Nguyen M, Zerbib D, Bernadou J, et al. In vitro inhibition of the Mycobacterium tuberculosis beta-ketoacyl-acyl carrier protein reductase MabA by isoniazid. Antimicrob Agents Chemother. 2004;48(1):242–9. Epub 2003/12/25. doi: 10.1128/AAC.48.1.242-249.2004. PubMed PMID: 14693546; PubMed Central PMCID: PMCPMC310174.

52. Samuel G, Reeves P. Biosynthesis of O-antigens: genes and pathways involved in nucleotide sugar precursor synthesis and O-antigen assembly. Carbohydr Res. 2003;338(23):2503–19. Epub 2003/12/13. doi: 10.1016/j.carres.2003.07.009. PubMed PMID: 14670712.

53. Alphey MS, Pirrie L, Torrie LS, Boulkeroua WA, Gardiner M, Sarkar A, et al. Allosteric competitive inhibitors of the glucose-1-phosphate thymidylyltransferase (RmlA) from Pseudomonas aeruginosa. ACS Chem Biol. 2013;8(2):387–96. Epub 2012/11/10. doi: 10.1021/cb300426u. PubMed PMID: 23138692.

54. Xiao G, Alphey MS, Tran F, Pirrie L, Milbeo P, Zhou Y, et al. Next generation Glucose-1-phosphate thymidylyltransferase (RmlA) inhibitors: An extended SAR study to direct future design. Bioorg Med Chem. 2021;50:116477. Epub 2021/11/11. doi: 10.1016/j.bmc.2021.116477. PubMed PMID: 34757294; PubMed Central PMCID: PMCPMC8613358.

55. Lewendon A, Coggins JR. 3-Phosphoshikimate 1-carboxyvinyltransferase from Escherichia coli. Methods in Enzymology. 1987;142:342–8.

56. Herrmann KM, Weaver LM. The Shikimate Pathway. Annu Rev Plant Physiol Plant Mol Biol. 1999;50:473–503. Epub 2004/03/12. doi: 10.1146/annurev.arplant.50.1.473. PubMed PMID: 15012217.

57. Stallings WC, Abdel-Meguid SS, Lim LW, Shieh HS, Dayringer HE, Leimgruber NK, et al. Structure and topological symmetry of the glyphosate target 5-enolpyruvylshikimate-3-phosphate synthase: a distinctive protein fold. Proc Natl Acad Sci U S A. 1991;88(11):5046–50. Epub 1991/06/01. doi: 10.1073/pnas.88.11.5046. PubMed PMID: 11607190; PubMed Central PMCID: PMCPMC51804.

58. Sterling T, Irwin JJ. ZINC 15--Ligand Discovery for Everyone. J Chem Inf Model. 2015;55(11):2324–37. Epub 20151109. doi: 10.1021/acs.jcim.5b00559. PubMed PMID: 26479676; PubMed Central PMCID: PMCPMC4658288.

59. Lipinski CA, Lombardo F, Dominy BW, Feeney PJ. Experimental and computational approaches to estimate solubility and permeability in drug discovery and development settings. Adv Drug Deliv Rev. 2001;46(1-3):3–26. Epub 2001/03/22. doi: S0169-409X(00)00129-0 [pii]. PubMed PMID: 11259830.

60. Morris GM, Huey R, Lindstrom W, Sanner MF, Belew RK, Goodsell DS, et al. AutoDock4 and AutoDockTools4: Automated docking with selective receptor flexibility. J Comput Chem. 2009;30(16):2785–91. Epub 2009/04/29. doi: 10.1002/jcc.21256. PubMed PMID: 19399780; PubMed Central PMCID: PMC2760638.

61. Moustakas DT, Lang PT, Pegg S, Pettersen E, Kuntz ID, Brooijmans N, et al. Development and validation of a modular, extensible docking program: DOCK 5. J Comput Aided Mol Des. 2006;20(10-11):601–19. Epub 2006/12/07. doi: 10.1007/s10822-006-9060-4. PubMed PMID: 17149653.

62. Guimaraes CR, Cardozo M. MM-GB/SA rescoring of docking poses in structure-based lead optimization. J Chem Inf Model. 2008;48(5):958–70. Epub 2008/04/22. doi: 10.1021/ci800004w. PubMed PMID: 18422307.

63. Deng Y, Roux B. Computation of binding free energy with molecular dynamics and grand canonical Monte Carlo simulations. J Chem Phys. 2008;128(11):115103. Epub 2008/03/26. doi: 10.1063/1.2842080. PubMed PMID: 18361618.

64. Siegel RL, Giaquinto AN, Jemal A. Cancer statistics, 2024. CA Cancer J Clin. 2024;74(1):12-49. Epub 20240117. doi: 10.3322/caac.21820. PubMed PMID: 38230766.

65. Xu L, Gordon R, Farmer R, Pattanayak A, Binkowski A, Huang X, et al. Precision therapeutic targeting of human cancer cell motility. Nat Commun. 2018;9(1):2454. Epub 20180622. doi: 10.1038/s41467-018-04465-5. PubMed PMID: 29934502; PubMed Central PMCID: PMCPMC6014988.

66. Wang LD, Wagers AJ. Dynamic niches in the origination and differentiation of haematopoietic stem cells. Nat Rev Mol Cell Biol. 2011;12(10):643–55. Epub 20110902. doi: 10.1038/nrm3184. PubMed PMID: 21886187; PubMed Central PMCID: PMCPMC4040463.

67. Hart L, Ogbonnaya A, Boykin K, Deyoung K, Bailey R, Heritage T, et al. Burden of chemotherapy-induced myelosuppression among patients with extensive-stage small cell lung cancer: A retrospective study from community oncology practices. Cancer Med. 2023;12(8):10020–30. Epub 20230331. doi: 10.1002/cam4.5738. PubMed PMID: 37000119; PubMed Central PMCID: PMCPMC10166910.

68. Epstein RS, Aapro MS, Basu Roy UK, Salimi T, Krenitsky J, Leone-Perkins ML, et al. Patient Burden and Real-World Management of Chemotherapy-Induced Myelosuppression: Results from an Online Survey of Patients with Solid Tumors. Adv Ther. 2020;37(8):3606–18. Epub 20200708. doi: 10.1007/s12325-020-01419-6. PubMed PMID: 32642965; PubMed Central PMCID: PMCPMC7340862.

69. Kurtin S. Myeloid toxicity of cancer treatment. J Adv Pract Oncol. 2012;3(4):209–24. PubMed PMID: 25031949; PubMed Central PMCID: PMCPMC4093344.

70. Purgato M, Barbui C. What is the WHO essential medicines list? Epidemiol Psychiatr Sci. 2012;21(4):343–5. Epub 20120730. doi: 10.1017/S204579601200039X. PubMed PMID: 22846155; PubMed Central PMCID: PMCPMC6998134.

71. Grem JL. 5-Fluorouracil: forty-plus and still ticking. A review of its preclinical and clinical development. Invest New Drugs. 2000;18(4):299–313. doi: 10.1023/a:1006416410198. PubMed PMID: 11081567.

72. Meredith JE, Jr., Fazeli B, Schwartz MA. The extracellular matrix as a cell survival factor. Mol Biol Cell. 1993;4(9):953–61. doi: 10.1091/mbc.4.9.953. PubMed PMID: 8257797; PubMed Central PMCID: PMCPMC275725.

73. Frisch SM, Francis H. Disruption of epithelial cell-matrix interactions induces apoptosis. J Cell Biol. 1994;124(4):619–26. doi: 10.1083/jcb.124.4.619. PubMed PMID: 8106557; PubMed Central PMCID: PMCPMC2119917.

74. Hu W, Ohue M. SpatialPPI: Three-dimensional space protein-protein interaction prediction with AlphaFold Multimer. Comput Struct Biotechnol J. 2024;23:1214–25. Epub 20240315. doi: 10.1016/j.csbj.2024.03.009. PubMed PMID: 38545599; PubMed Central PMCID: PMCPMC10966450.

75. Gorostiola Gonzalez M, Janssen APA, AP IJ, Heitman LH, van Westen GJP. Oncological drug discovery: AI meets structure-based computational research. Drug Discov Today. 2022;27(6):1661–70. Epub 20220314. doi: 10.1016/j.drudis.2022.03.005. PubMed PMID: 35301149.

76. Arul Murugan N, Ruba Priya G, Narahari Sastry G, Markidis S. Artificial intelligence in virtual screening: Models versus experiments. Drug Discov Today. 2022;27(7):1913–23. Epub 20220518. doi: 10.1016/j.drudis.2022.05.013. PubMed PMID: 35597513.

77. Gedgaudas M, Baronas D, Kazlauskas E, Petrauskas V, Matulis D. Thermott: A comprehensive online tool for protein-ligand binding constant determination. Drug Discov Today. 2022;27(8):2076–9. Epub 20220514. doi: 10.1016/j.drudis.2022.05.008. PubMed PMID: 35577233.

78. Kunimoto R, Bajorath J, Aoki K. From traditional to data-driven medicinal chemistry: A case study. Drug Discov Today. 2022;27(8):2065–70. Epub 20220420. doi: 10.1016/j.drudis.2022.04.017. PubMed PMID: 35452790.

79. Vermeulen I, Isin EM, Barton P, Cillero-Pastor B, Heeren RMA. Multimodal molecular imaging in drug discovery and development. Drug Discov Today. 2022;27(8):2086–99. Epub 20220413. doi: 10.1016/j.drudis.2022.04.009. PubMed PMID: 35429672.

80. Paoli P, Giannoni E, Chiarugi P. Anoikis molecular pathways and its role in cancer progression. Biochim Biophys Acta. 2013;1833(12):3481–98. Epub 20130702. doi: 10.1016/j.bbamcr.2013.06.026. PubMed PMID: 23830918.

81. Neuendorf HM, Simmons JL, Boyle GM. Therapeutic targeting of anoikis resistance in cutaneous melanoma metastasis. Front Cell Dev Biol. 2023;11:1183328. Epub 20230426. doi: 10.3389/fcell.2023.1183328. PubMed PMID: 37181747; PubMed Central PMCID: PMCPMC10169659.

82. AI’s potential to accelerate drug discovery needs a reality check. Nature. 2023;622(7982):217. doi: 10.1038/d41586-023-03172-6. PubMed PMID: 37817040.

83. Liu YQ, Kyle E, Patel S, Housseau F, Hakim F, Lieberman R, et al. Prostate cancer chemoprevention agents exhibit selective activity against early stage prostate cancer cells. Prostate cancer and prostatic diseases. 2001;4(2):81–91. Epub 2002/12/24. doi: 10.1038/sj.pcan.4500506. PubMed PMID: 12497043.

84. Xu L, Ding Y, Catalona WJ, Yang XJ, Anderson WF, Jovanovic B, et al. MEK4 function, genistein treatment, and invasion of human prostate cancer cells. J Natl Cancer Inst. 2009;101(16):1141–55. Epub 20090728. doi: 10.1093/jnci/djp227. PubMed PMID: 19638505; PubMed Central PMCID: PMCPMC2728746.

85. Zhang L, Pattanayak A, Li W, Ko HK, Fowler G, Gordon R, et al. A Multifunctional Therapy Approach for Cancer: Targeting Raf1-Mediated Inhibition of Cell Motility, Growth, and Interaction with the Microenvironment. Mol Cancer Ther. 2020;19(1):39–51. Epub 20191003. doi: 10.1158/1535-7163.MCT-19-0222. PubMed PMID: 31582531.

86. Westbrook J, Feng Z, Chen L, Yang H, Berman HM. The Protein Data Bank and structural genomics. Nucleic Acids Res. 2003;31(1):489–91. PubMed PMID: 12520059.

87. Huang X, Chen S, Xu L, Liu Y, Deb DK, Platanias LC, et al. Genistein inhibits p38 map kinase activation, matrix metalloproteinase type 2, and cell invasion in human prostate epithelial cells. Cancer Res. 2005;65(8):3470–8. doi: 10.1158/0008-5472.CAN-04-2807. PubMed PMID: 15833883.

